# Rapid Evolution of Knockdown Resistance Haplotypes in Response to Pyrethroid Selection in *Aedes aegypti*

**DOI:** 10.1101/2021.04.02.438212

**Authors:** Jennifer Baltzegar, Michael Vella, Christian Gunning, Gissella Vasquez, Helvio Astete, Fred Stell, Michael Fisher, Thomas W. Scott, Audrey Lenhart, Alun L. Lloyd, Amy Morrison, Fred Gould

**Author notes:** **Data for this study are available at:** to be completed after manuscript is accepted for publication. **Disclaimer** The views expressed in this manuscript are those of the authors and do not necessarily reflect the official policy or position of the Department of the Navy, Department of Defense, the Centers for Disease Control and Prevention, nor the U.S. Government. **Copyright statement** Some authors of this manuscript are military service members and employees of the U.S. Government. This work was prepared as part of their official duties. Title 17 U.S.C. §105 provides that “Copyright protection under this Title is not available for any work of the United States Government”. Title 17 U.S.C. §101 defines a U.S. Government work as a work prepared by a military service member or employee of the U.S. Government as part of that person’s official duties.

## Abstract

This study describes the evolution of *knockdown resistance (kdr)* haplotypes in *Aedes aegypti* in response to pyrethroid insecticide use over the course of 18 years in Iquitos, Peru. Based on the duration and intensiveness of sampling (∼10,000 samples), this is the most thorough study of kdr population genetics in *Ae. aegypti* to date within a city. We provide evidence for the direct connection between programmatic citywide pyrethroid spraying and the increase in frequency of specific *kdr* haplotypes by identifying two evolutionary events in the population. The relatively high selection coefficients, even under infrequent insecticide pressure, emphasizes how quickly populations can evolve. The observed rapid increase in frequency of resistance alleles might have been aided by the incomplete dominance of resistance-conferring alleles over corresponding susceptibility alleles. In addition to dramatic temporal shifts, spatial suppression experiments reveal that genetic heterogeneity existed not only at the citywide scale, but also on a very fine scale within the city.

## Introduction

The *Aedes aegypti* (L.) mosquito vectors yellow fever, dengue, Zika, and chikungunya viruses. Of these viruses, dengue virus (DENV) is the most impactful, infecting almost 400 million people annually (Bhatt *et al.,* 2013) and causing an estimated 96 million symptomatic cases and 40,500 deaths every year (GBD 2017 Causes of Death Collaborators, 2018). While a dengue vaccine is approved in certain parts of the world, the World Health Organization (WHO) recommends that access be restricted by age and prior dengue infection before it can be safely administered (Thomas and Yoon, 2019). Dengue control has also been attempted by releasing laboratory-manipulated mosquitoes to either suppress populations or to render them unable to transmit DENV to humans (e.g., Hoffmann *et al.,* 2011; Harris *et al.,* 2012; Lacroix *et al.,* 2012; Carvalho *et al.,* 2015; Nguyen *et al.,* 2015). The most common approach to reducing dengue disease incidence, however, continues to be vector control efforts such as space and residual surface spraying of chemical insecticides.

Because pyrethroids exhibit low mammalian toxicity, they have become a preferred tool for *Ae. aegypti* population control. Pyrethroids are highly effective against *Ae. aegypti* because they target the voltage-gated sodium channel protein (VGSC), which is coded for by a single copy gene within the *Ae. aegypti* genome (Dong *et al.,* 2014). However, resistance to pyrethroids in *Ae. aegypti* is becoming widespread throughout the world (e.g., Smith *et al.,* 2016; Bharati and Saha, 2018; Kamgang *et al.,* 2017; Flores-Suarez *et al.,* 2016; Li *et al.,* 2015; Nguyen *et al.,* 2018).

Considerable genetic research has been conducted on single nucleotide mutations in the *VGSC* gene that contribute to phenotypic pyrethroid resistance in *Ae. aegypti* (reviewed in Du *et al.,* 2016). In Central and South America, pyrethroid resistance in *Ae. aegypti* has been associated with single nucleotide mutations that cause amino acid changes from valine (V) to isoleucine (I) at position 1016 (V1016I), and from phenylalanine (F) to cysteine (C) at position 1534 (F1534C), (numbered according to homology in the house fly, *Musca domestica*) in the VGSC protein (Tbl. 1) (Linss *et al.,* 2014; Deming *et al.,* 2016). Another common single nucleotide mutation in Brazil and Mexico that is associated with pyrethroid resistance is the valine (V) to leucine (L) amino acid substitution at locus 410 (Haddi *et al.,* 2017; Saveedra-Rodriguez *et al.,* 2018). Different non-synonymous mutations in the *VGSC* gene are more common in other parts of the world. For example, the single nucleotide mutation resulting in the V1016G substitution is very common in Asia (Li *et al.,* 2015) and has only recently been reported in the Western Hemisphere (Murcia *et al.,* 2019).

Since the mutations causing V1016I and F1534C are both within the *VGSC* gene, they are expected to be in linkage disequilibrium when polymorphic (Hartl, 2000). Haplotypes that contain both the Cys1534 allele and the wild-type Val1016 allele are common in wild populations (Linss *et al*., 2014, Deming *et al*., 2016). In contrast, the Ile1016 allele is rarely found on the same chromosome as the wild type Phe1534 allele (Linss *et al.,* 2014). Vera-Maloof *et al*. (2015) suggest that the Cys1534 allele may be required to support the presence of the Ile1016 allele, yet these resistance alleles do not always rise in frequency together. In Brazil, the frequency of both resistance alleles, Ile1016 and Cys1534, increased over time at all sites sampled, but the ratio of different haplotypes of the gene with one or both nucleotide changes varied geographically (Linss *et al.,* 2014). More recently, *Ae. aegypti kdr* haplotype analysis has confirmed that the Val1016/Cys1534 haplotype is found worldwide, while the Ile1016/Cys1534 haplotype is found only in the Americas (Cosme *et al.,* 2020; Fan *et al.,* 2020). This implies that both widely-dispersed and geographically constrained haplotypes contribute to the wide-spread pyrethroid resistance observed in *Ae. aegypti*.

Although specific resistance haplotypes are commonly observed in *Ae. aegypti* in the Western Hemisphere, results from two studies in Merida, Mexico indicate that heterogeneity in frequencies of common resistance haplotypes can exist at the scale of a city block (Deming *et al.,* 2016). High spatial heterogeneity means that fine-scale spatial variability is not always observable when assessing insecticide resistance at city or region-wide scales (Grossman *et al.,* 2019). These two studies are important for understanding the scale of geographic variability in *Ae. aegypti* insecticide resistance and the role that limited mobility of this species plays in this evolutionary process. These studies were not, however, able to determine if the heterogeneity in resistance is due to spatial variation in insecticide pressure or to variation in initial resistance allele frequencies. A better understanding of fine-scale spatial structure of resistance genetics can inform response to control measures, thus preserving efficacy of important insecticides and aiding in the mathematical modeling of new insect control technologies.

Iquitos, Peru is a well-established study site for *Ae. aegypti* in the Western Hemisphere with a long history of research and sampling that can help elucidate the factors contributing to pyrethroid resistance evolution in this mosquito. Iquitos is located in the northeastern Peruvian Amazon and is the largest city in the world without major connecting roads. The population was estimated to be ∼437,300 in 2015 (Instituto Nacional de Estadistica e Informatica, 2012). *Ae. aegypti* were initially eradicated from Peru in 1958, but they reinvaded and were detected again in 1984 (Phillips *et al*., 1992). In anticipation of dengue moving into areas with *Ae. aegypti* populations, the U.S. Naval Medical Research Unit No. 6 (NAMRU-6) established a field office in Iquitos in the late 1980s to monitor for Aedes-transmitted viruses. And, as local DENV transmission has occurred since the early 1990s, repeated citywide insecticide spraying and long-term epidemiological monitoring efforts have been carried out to control *Ae. aegypti* populations to reduce disease (Morrison *et al*., 2010; Stoddard *et al*., 2014). Dengue-1 was detected in Iquitos in 1990 (Phillips *et al*., 1992) and dengue-2 was confirmed in 1995 (Watts *et al*., 1999). Continuous hospital and outpatient clinical surveillance began in 1993 (Watts *et al*., 1999), and household vector surveillance began in 1998 (Morrison *et al*., 2004). The household vector surveillance collection spans the years before, during, and after pyrethroid use in the city, making Iquitos an excellent location to explore the natural evolution of pyrethroid resistance in *Ae. aegypti*.

Here, we examined the temporal and spatial patterns of *kdr* allele frequencies in *Ae. aegypti* in Iquitos, Peru. We analyze the timing and rate at which haplotypes containing the two most commonly observed *kdr* mutations in Central and South America, V1016I and F1534C, emerged in the city over the course of 18 years (∼180 - 216 generations). We assess the importance that spatial heterogeneity and dominance play in this dynamic and investigate the strength of selection on these two resistance alleles during periods of pyrethroid selection.

## Materials and Methods

### Entomological Surveys

#### Control History

During the past two decades, researchers have preserved mosquito specimens collected throughout the city of Iquitos (e.g. Cavany *et al*., 2020; Cromwell *et al*., 2017; Getis *et al.,* 2003; Gunning *et al.,* 2018; LaCon *et al.,* 2014; Lenhart *et al*., 2020; Morrison *et al.,* 2004a; Morrison *et al.,* 2004b; Morrison *et al.,* 2006; Morrison *et al.,* 2008; Reiner *et al*., 2019; Schneider *et al*., 2004; Tun-Lin *et al*., 2009). Prior to 2002, no citywide insecticide spraying occurred. Multiple sub-classes of pyrethroid insecticides were sprayed by the Iquitos ministry inside homes from 2002 – 2014. These sub-classes included deltamethrin, cypermethrin, alpha-cypermethrin, lambda-cyhalothrin, and alpha-cypermethrin + pyriproxyfen (Suppl. Tbl. 1). It was uncommon for residents to spray their own homes (A. Morrison, personal communication). During 2014, pyrethroids were discontinued in favor of malathion due to the development of phenotypic pyrethroid resistance (Gunning *et al.,* 2018). Over the 18-year period, spraying occurred at an average of 3.25 treatments per year, with spraying occurring within a one-month period in most years. Because the applications were Ultra Low Volume space sprays with very short persistence, most generations of the mosquitoes in any year were not exposed to insecticide.

#### Temporal Collections

*Aedes aegypti* were collected and stored at −80°C by NAMRU-6 and University of California - Davis personnel since the late 1990s. Specimens dating back to the year 2000 are still available for study. Mosquitoes were collected by backpack aspirator (Clark *et al.,* 1994) prior to June 2009 and by Prokopack Aspirator following June 2009 (Vasquez-Prokopec *et al*., 2009; Reiner *et al*., 2019). Each mosquito in the repository was identified to species, sex, collection date, and collection site. Each collection site, typically an individual household, was associated with GPS coordinates (Fig. 1).

**Figure 1.**
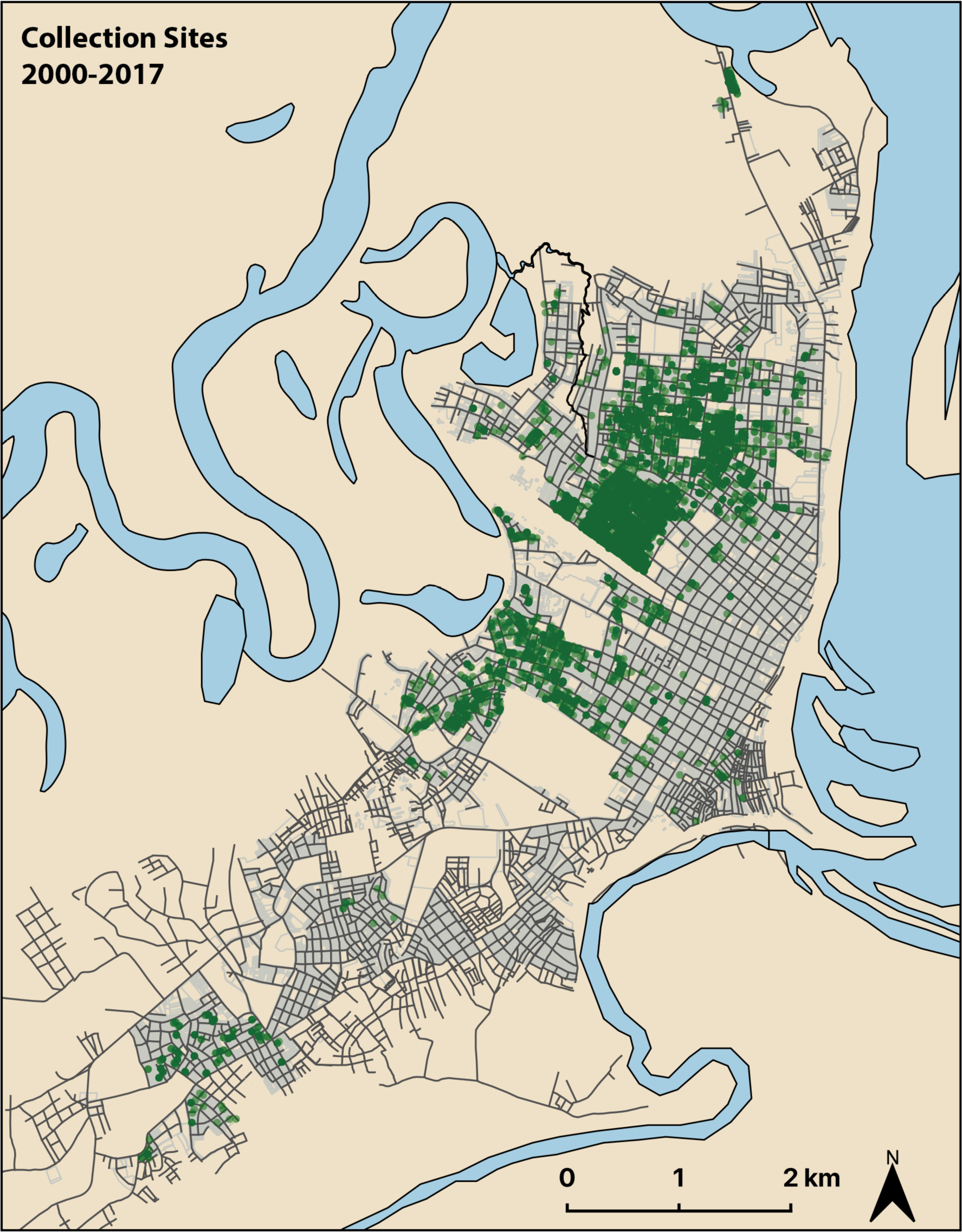
Map of Iquitos, Peru showing collection sites for *Ae. aegypti* collected from 2000 - 2017. Black lines represent roadways. Collection sites are represented by green circles. Darker green indicates more sampling.

#### Spatial Collections

Intense suppression experiments based on pyrethroid spraying were conducted in 2013 and 2014 (Gunning *et al.,* 2018) to test the predictions of a detailed *Ae. aegypti* population dynamics model (Magori *et al.,* 2009). In brief, two areas of the city were identified as having relatively high densities of *Ae. aegypti* and were configured spatially in a way that allowed for a central spray sector with an outer buffer sector to act as an experimental control region (Fig. 2). The 2013 study site covered approximately 750 m x 450 m and contained 1,163 houses. Baseline samples were collected in January 2013 and systematic sampling for this study began on 22 April 2013 and continued for 16 calendar weeks until 8 August 2013. From 29 April 2013 to 3 June 2013, six weekly non-residual, indoor ultra-low volume (ULV) cypermethrin treatments were applied in the treatment sector. The 2014 study site was larger and covered an approximate 600 m x 600 m area and contained 2,166 houses. Systematic sampling was conducted over a longer period of 44 calendar weeks. ULV spraying of cypermethrin was performed from 28 April 2014 through 2 June 2014 in a similar manner as in the 2013 study. In addition to the study spray in 2014, a citywide spray was conducted in response to a dengue outbreak in February 2014, where homes in both the experimental and buffer sectors were sprayed with pyrethroids. Throughout both suppression experiments, mosquitoes were collected and stored as described above.

**Figure 2.**
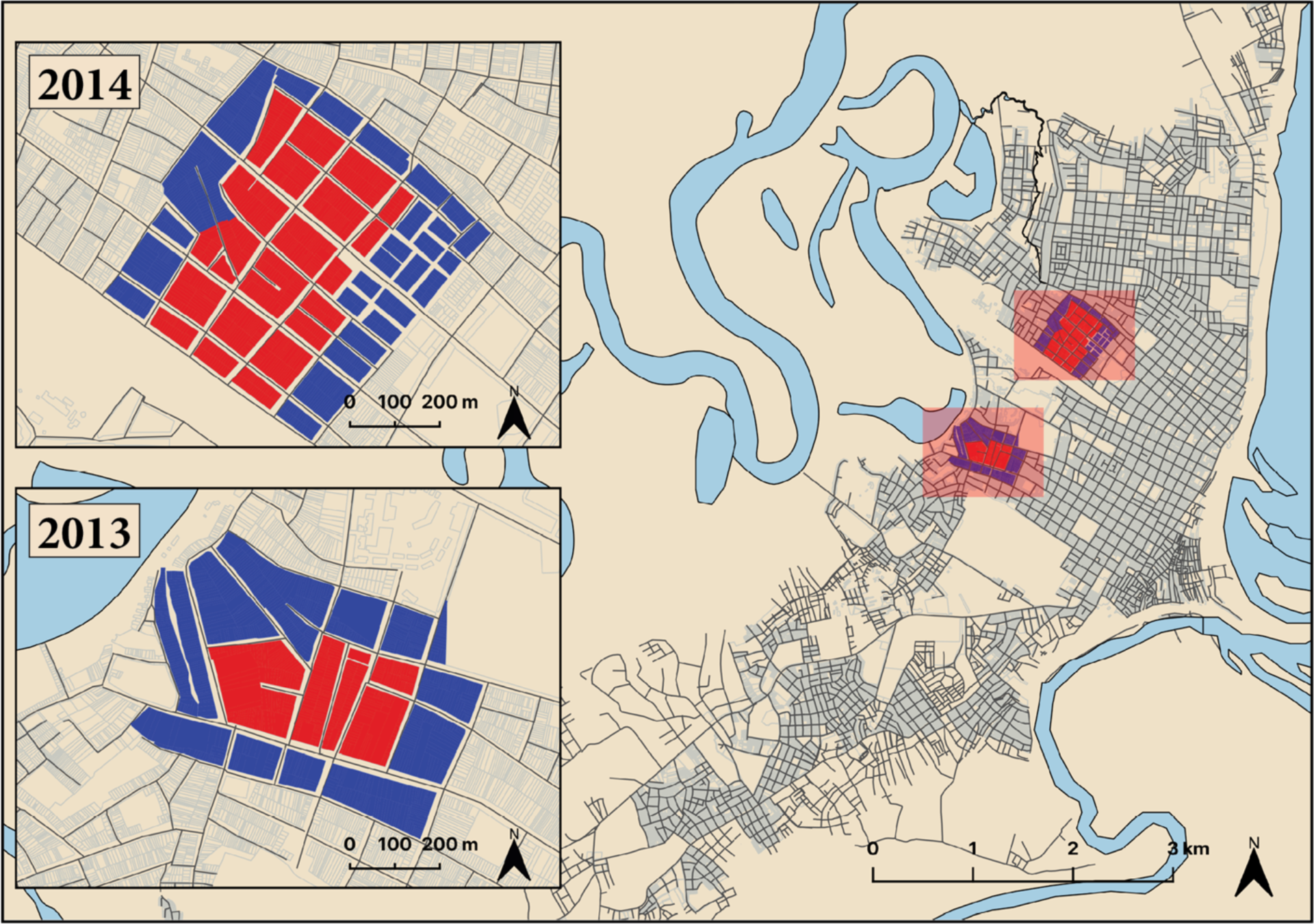
Maps of the 2013 and 2014 suppression experiments showing the buffer (blue) and spray (red) zones from where *Ae. aegypti* were collected. Many houses, often sharing walls, form blocks that are separated by roadways (black lines).

### DNA Extractions and Quantification

Male mosquitoes were chosen for genetic analysis throughout this study because female mosquitoes were typically the focus of virology and epidemiological studies and, therefore more males were available in the repository. Using females would also have brought the risk of genomic contamination from male mosquitoes (via insemination) and from humans (via blood feeding). Males are expected to share similar allele frequencies with females because the *VGSC* is not sex-linked.

Whole mosquitoes were transferred from Iquitos, Peru to Raleigh, North Carolina, USA with permits from both Peruvian and US authorities. Samples were stored at −80°C prior to genomic DNA (gDNA) isolation and at −20°C after gDNA isolation. Genomic DNA was extracted from whole male *Ae. aegypti* by one of two methods: Qiagen DNeasy blood and tissue kit (cat: 69582) or Canadian Center for DNA Barcoding protocol. In brief, for the Qiagen DNeasy kit protocol, whole male mosquitoes were homogenized and incubated in lysis buffer and proteinase K overnight at 55°C. Following incubation and removal of chitinous material, RNase A treatment was performed to remove RNA contamination for both isolation methods. Then, the standard Qiagen protocol of washes was followed. Final samples were eluted two times in 150 µl warm dH_2_O (Invitrogen Cat #: 10977-015). A modified Canadian Center for DNA Barcoding (2020) protocol was also used for some mosquito DNA isolations to reduce costs while maintaining quality genomic DNA extractions. Samples were homogenized, incubated, and RNase A treated as described above before the lysate was passed through the filter of an AcroPrep™ PALL2 plate (Cat #: PALL 5053) to bind the gDNA. The filter was washed with Protein Wash Buffer to remove remaining proteins and then washed with cold Wash Buffer to remove additional contaminates. The filter was allowed to dry to ensure that no ethanol remained to interfere with DNA yield. Finally, two washes of 75 µl warm dH_2_O (Invitrogen Cat #: 10977-015) were performed to elute a final volume of 175 µl gDNA.

Quantification of gDNA was performed using a Quant-iT PicoGreen dsDNA assay (ThermoFisher Scientific - P11496) and samples were read on a Synergy H1 Hybrid Plate Reader (BioTek Instruments, Inc.) in the Genomic Sciences Laboratory at North Carolina State University (GSL).

### Genotyping

Allele-specific quantitative PCR and melting curve analysis (AS-PCR) was used to genotype all mosquitoes in duplicate for each of the mutations most commonly found in Central and South America (V1016I and F1534C). If the two reactions were not scored identically, the sample was discarded from further analysis. Mismatches were rare and typically due to non-amplification of a sample or because certain criteria for scoring were not met (i.e., melting peak did not cross threshold). A smaller number (n=92) of individuals were additionally genotyped at the V410L locus to verify the strong linkage disequilibrium that has been previously reported between it and locus V1016I (Saavedra-Rodriguez *et al.,* 2018). Each mosquito genotyped at the V410L locus was also genotyped twice to ensure accuracy.

#### Genotyping of V1016I

AS-PCR for the V1016I locus was based on the method reported by Saavedra-Rodriguez *et al*. (2007) and modifications to the I1016R primer made by the Entomology Branch at the Centers for Disease Control and Prevention (CDC), Atlanta, USA (A. Lenhart, personal communication). The PCR volume was reduced to 10 µl per reaction and contained 2.5 µl of dH_2_O, 0.5 µl of each primer at 10 µM (V1016F, I1016F, I1016R), 5 µl of PerfeCTa SYBR Green Supermix (Quanta – 95054-02K), and 1 µl of template. The primer sequences for V1016F and I1016F used are reported in Saavedra-Rodriquez *et al*. (2007), but the primer sequence for I1016R was modified to: 5’ - TGA TGA ACC SGA ATT GGA CAA AAG C – 3’ (CDC, personal communication). Samples were genotyped on a BioRad CFX384 Real-Time PCR machine in the GSL, with the following thermal conditions: step 1 - 95°C for 3 minutes, step 2 – 95°C for 10 seconds, step 3 – 60°C for 10 seconds, step 4 – 72°C for 10 seconds, step 5 - Go to step 2, 39 times, step 6 – 95°C for 10 seconds, step 7 - Melting Curve 65°C - 95°C, increment 0.2°C per 10 sec plus a plate read.

#### Genotyping of F1534C

AS-PCR for the F1534C locus was performed following the method reported by Yanola *et al*. (2011) with the following modifications. The PCR volume was reduced to a total of 10 µl per reaction and contained 5 µl dH_2_O, 0.2 µl C1534F primer, 0.4 µl F1534F primer, 0.4 µl F1534R primer, 3.0 µl of PerfeCTa SYBR Green Supermix (Quanta – 95054-02K), and 1 µl of template. Samples were genotyped on a BioRad CFX384 Real-Time PCR machine in the GSL, with the following thermal conditions: step 1 - 95°C for 2 minutes, step 2 – 95°C for 30 seconds, step 3 – 60°C for 30 seconds, step 4 – 72°C for 30 seconds, step 5 - Go to step 2, 34 times, step 6 – 72°C for 2 minutes, step 7 - Melting Curve 65°C - 95°C, increment 0.2°C per 10 sec plus a plate read.

#### Genotyping of V410L

The AS-PCR for the V410L locus was based on a protocol developed by K. Saavedra and shared via the Entomology Branch, CDC (A. Lenhart, personal communication). The total volume for each reaction was reduced to 10 µl: 3.8 µl dH_2_O, 0.05 µl Val410 primer (50 µM) 5’ – GCG GGC AGG GCG GCG GGG GCG GGG CCA TCT TCT TGG GTT CGT TCT ACC GTG – 3’, 0.05 µl Leu410 primer (50 µM) 5’ – GCG GGC ATC TTC TTG GGT TCG TTC TAC CAT T – 3’, 0.1 µl Rev410 primer (50 µM) 5’ – TTC TTC CTC GGC GGC CTC TT – 3’, 5.0 µl PerfeCTa SYBR Green SuperMix (Quanta – 95054-02K), and 1 µl template. Thermal conditions were performed on the BioRad CFX384 Real-Time PCR machine in the GSL, with the following thermal conditions: 95°C for 3:00, 40 cycles of (95°C for 0:10, 60°C for 0:10, 72°C for 0:30), 95°C for 0:10, melting curve 65°C – 95°C increasing in increments of 0.2°C per 10 sec plus a plate read.

#### Analysis for AS-PCR

Melt Curve peak calls were determined using the CFX Maestro Software (Bio-Rad - 12004110), verified by eye, and exported to a customized C++ script to quickly convert melt curve peak calls to genotypes for each sample. Melt curve genotypes were then read into a customized R script (R Core Team 2019) for allele frequency determination and other statistical analysis.

### Statistical Analysis

#### Pairwise Linkage Disequilibrium between SNP markers

Linkage disequilibrium (LD) was calculated between V1016I and V410L for years after emergence using the ‘genetics’ package in R (v. 3.5.0) (Warnes *et al.,* 2019). Pairwise LD was neither calculated between V1016I and F1534C nor F1534C and V410L because Ile1016 and Leu410 did not appear in the population until Cys1534 was near fixation (see results).

#### Imputation of Haplotypes

For individual mosquitoes with at least one homozygous genotype at either locus, haplotypes could be known with confidence. For individual mosquitoes that were heterozygous at either the 1534 and 1016 loci, imputed haplotype frequencies were used for analysis as there is a high degree of physical linkage between the SNP loci examined for this study. To impute the haplotypes, we implemented a custom script in R by first creating a matrix of haplotype probabilities given specific SNP genotypes and considering the assumptions about the existence of the Ile1016/wt1534 haplotype. We then multiplied the genotype count matrix by the probability matrix and summed across the nine genotypes to achieve an estimate of individuals with a given haplotype per year. Finally, haplotype frequencies were calculated from this estimate. For the prior estimate we assume 1) Mendelian inheritance at both loci and 2) that the resistance allele Ile1016 never occurs with the susceptible allele Phe1534. While this assumption may introduce small errors in the imputation, it is reasonable because the frequency of the Cys1534 allele reaches fixation in the sampled population prior to the emergence of the Ile1016 allele (Fig. 3). We observed only 3 out of 9,882 individuals (0.03%) which had to have contained this haplotype (data not shown). Thus, it is likely that few Ile1016/Phe1534 haplotypes existed in the Iquitos population during this study period. Previous investigators have rarely reported instances of this haplotype occurring in Central and South America (Linss *et al.,* 2014; Vera-Maloof *et al.,* 2015; Deming *et al.,* 2016; Grossman *et al.,* 2019). Vera-Maloof *et al*. (2015) reported the largest frequency (0.09) of this haplotype, but the frequency decreased in the same area in subsequent collections and they concluded that it must have low fitness, even in the presence of pyrethroids.

**Figure 3.**
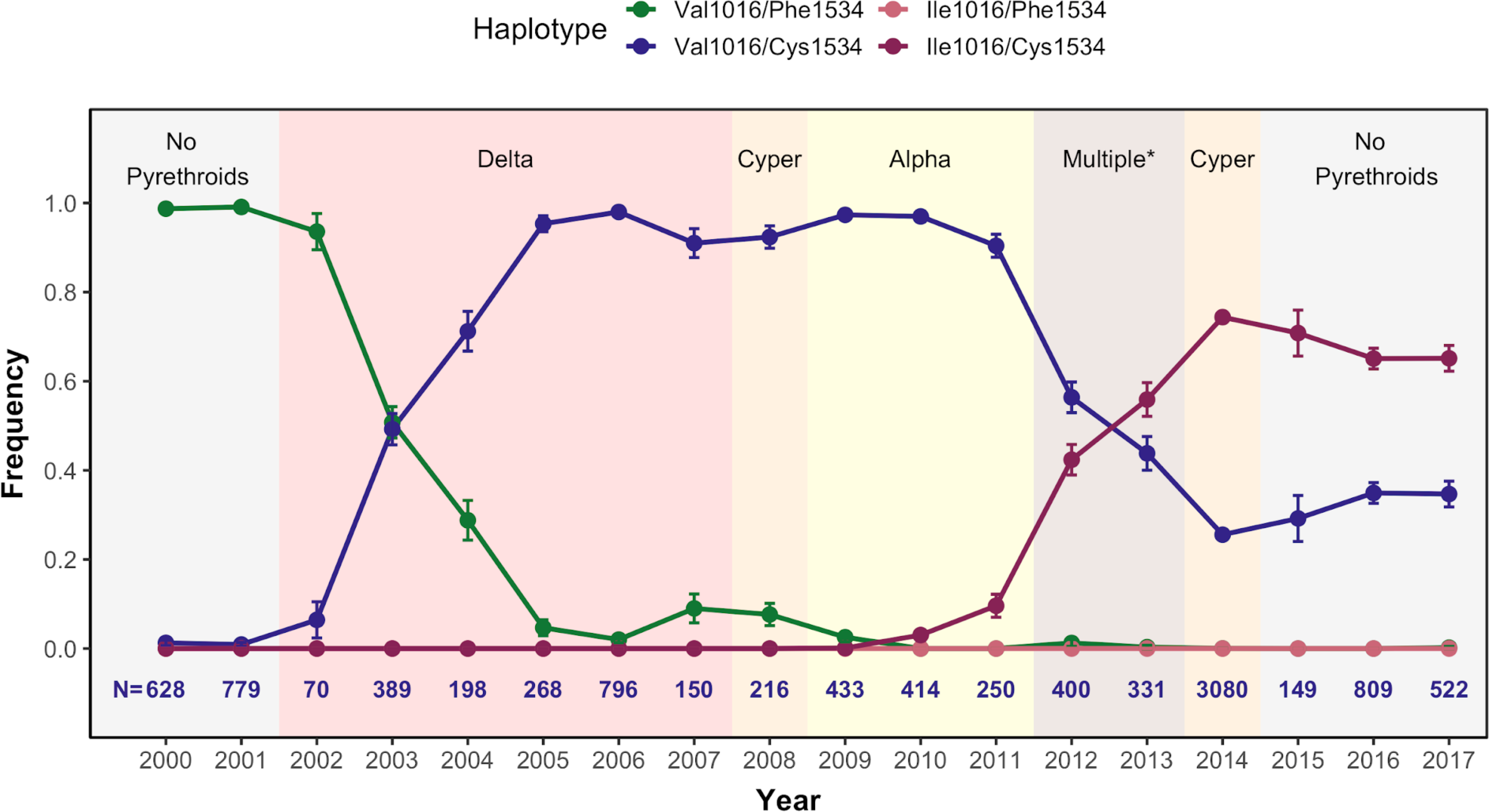
*kdr* haplotype frequencies from 2000 - 2017. Shaded areas represent periods when pyrethroids were used or not. The specific type of pyrethroid is abbreviated in the shaded area (i.e., Delta = deltamethrin, Cyper = cypermethrin, and Alpha = alpha-cypermethrin). Multiple* represents a period from 2012 – 2013 where multiple chemistries were used, including cypermethrin, alpha-cypermethrin, lambda-cyhalothrin, and alpha-cypermethrin+pyriproxyfen. Number of individuals genotyped per year are shown below y = 0.0. Error Bars = 95% CI.

#### Statistical Analyses

For each year, we conducted a logistic regression to compute confidence intervals on monthly resistance allele frequencies in the spray and buffer zones. We then used these regression models to compute contrast ratios between zones and their corresponding p-values in R via the ‘emmeans’ package (Lenth, 2020). For specific months (October 2002 - July 2004) during the increase of the Val1016/Cys1534 haplotype, we tested for Hardy-Weinberg Equilibrium (HWE) using the *‘genetics’* package in R (v. 3.5.0) and applied a Bonferroni correction (Warnes *et al.,* 2019).

### Parameter Estimation of Selection Coefficient and Dominance

#### Parameter Estimation using WFABC

The selection coefficient (s) and dominance (h) of the resistance allele was estimated at each locus in the population for the periods when each resistance allele was increasing in frequency by following the Wright-Fisher approximate Bayesian computation (WFABC) method for temporally sampled data (Foll *et al.,* 2014). Data were input into the model based on the month of collection and the selection period was set as April 2010 – June 2014 and October 2002 – December 2009 for locus 1016 and 1534, respectively. The WFABC model can first estimate the effective population size (N_e_) to then use as a prior for determining the selection coefficient, but it assumes a genome wide sampling of mostly neutral loci. Since two non-neutral loci were genotyped, we set 2N_e_ equal to 2 times 500 based on previous research in *Ae. aegypti* (Saarman *et al.,* 2017). Saarman *et al*. (2017) estimated a wide window for N_e_ (25 - 3000), but found that most populations averaged between 400 - 600 for this parameter. We set min_s = −1, max_s = 0, min_h = 0, and max_h = 1. In the WFABC model, genotype fitness assumes a selection benefit to the susceptible allele. Since the susceptible allele is decreasing during the sampled period, we expect a negative selection coefficient. The additional modeling that we implement, described below, uses relative genotype fitness formulas that assume a selection cost to the susceptible allele. Therefore, we report the absolute value of the WFABC selection parameters to present them in terms of cost to the susceptible allele and to more easily compare results between the WFABC and genotype-frequency models. In our results, when dominance is zero, the heterozygote behaves exactly like the resistant homozygote and the resistance allele is considered dominant. Whereas, when dominance is one, the heterozygote behaves exactly like the susceptible homozygote and the resistance allele is considered recessive.

#### Parameter Estimation using a Genotype-Frequency Model

We use a genotype-frequency model to evaluate the spread of the resistance allele at the 1534 locus of the *VGSC*. We assumed the population was well-mixed with random mating and that there were discrete (non-overlapping) generations of mosquitoes, each lasting one month. We also assumed the population was isolated, without immigration or emigration of mosquitoes.

The full population genetics model (supplemental methods 1) and sampling distribution form a Hidden Markov Model (HMM). We conduct Bayesian inference on the HMM using particle Markov chain Monte Carlo (pMCMC). We implement pMCMC with a multivariate normal proposal distribution and adaptive Metropolis-Hastings acceptance in R (R Core Team 2019) using the package nimble (de Valpine *et al.,* 2017). We used 1000 particles for the particle filter. For parameters in the range [0,1] (*s*, *h*, and the initial frequency, *R*_0_, of the resistance allele), we use uninformative priors of Beta(1,1). For the overdispersion parameter, *A*, we use the uninformative prior *A* ∼ *Gamma*(0.01,0.01). To improve the time to convergence, we initialize the parameters using their maximum likelihood estimates. The output of pMCMC is the joint posterior distribution of the parameters along with samples of full time series of genotype frequencies, which we used to construct 95% credible intervals of genotype frequencies.

## Results

### Entomological Collections

#### Temporal Collections

The number of mosquitoes (Tbl. 1) and sampling areas (Fig. 1) vary across years and depend on a number of factors, including where *Ae. aegypti* surveys were conducted in any given year and availability of samples. The number of mosquitoes available for use per year ranged from less than 70 in 2002 to 3,080 in 2014. A total of 9,882 individuals were analyzed in this study.

#### Spatial Collections

Samples collected in 2013 and 2014 were used to assess spatial variations in allele frequencies because special attention had been paid to the exact location of houses where samples were collected, and the insecticide treatment history of the house (Fig. 2). Details can be found in Gunning *et al*. (2018). Of the total samples collected in 2013 and 2014, 372 males (collected in 2013) and 2,178 males (collected in 2014) were analyzed from the area where the suppression experiments were conducted (Tbl. 1).

**Table 1.** The number of mosquitoes that were genotyped and used for analysis in this study. Total males analyzed is the number of males that were genotyped at both the V1016I and F1534C loci and used in the haplotype analysis. Buffer and spray zones correspond to the subset of the total males analyzed that were collected in each area during the 2013 and 2014 suppression experiments. The discrepancy between the totals is due to mosquitoes collected outside of the study area in that year. The V410L genotyped column indicates the subset of total males that were genotyped at the 410 locus in each year.

| Year | Total<br>Analyzed | Buffer | Spray | V410L<br>Genotyped |
| --- | --- | --- | --- | --- |
| 2000 | 628 |  |  | 96 |
| 2001 | 779 |  |  | 103 |
| 2002 | 70 |  |  | 0 |
| 2003 | 389 |  |  | 94 |
| 2004 | 198 |  |  | 92 |
| 2005 | 268 |  |  | 95 |
| 2006 | 796 |  |  | 95 |
| 2007 | 150 |  |  | 6 |
| 2008 | 216 |  |  | 44 |
| 2009 | 433 |  |  | 0 |
| 2010 | 414 |  |  | 98 |
| 2011 | 250 |  |  | 95 |
| 2012 | 400 |  |  | 92 |
| 2013 | 331 | 252 | 120 | 95 |
| 2014 | 3080 | 1001 | 1177 | 70 |
| 2015 | 149 |  |  | 95 |
| 2016 | 809 |  |  | 94 |
| 2017 | 522 |  |  | 92 |

### Linkage Disequilibrium Results

Linkage disequilibrium between alleles at loci coding for V1016I and V410L was calculated for each year from 2010 to 2017 (Tbl. 2). For all timepoints, LD was significant with a Χ^2^ p-value <0.001 and an R^2^ range of 0.666 to 0.953. The data indicate that alleles at the two loci were in strong, but not perfect, linkage disequilibrium. Because the R^2^ values were high for alleles at loci V410L and V1016I, the genotype at locus 1016 were used to predict the genotype at locus 410.

**Table 2.**
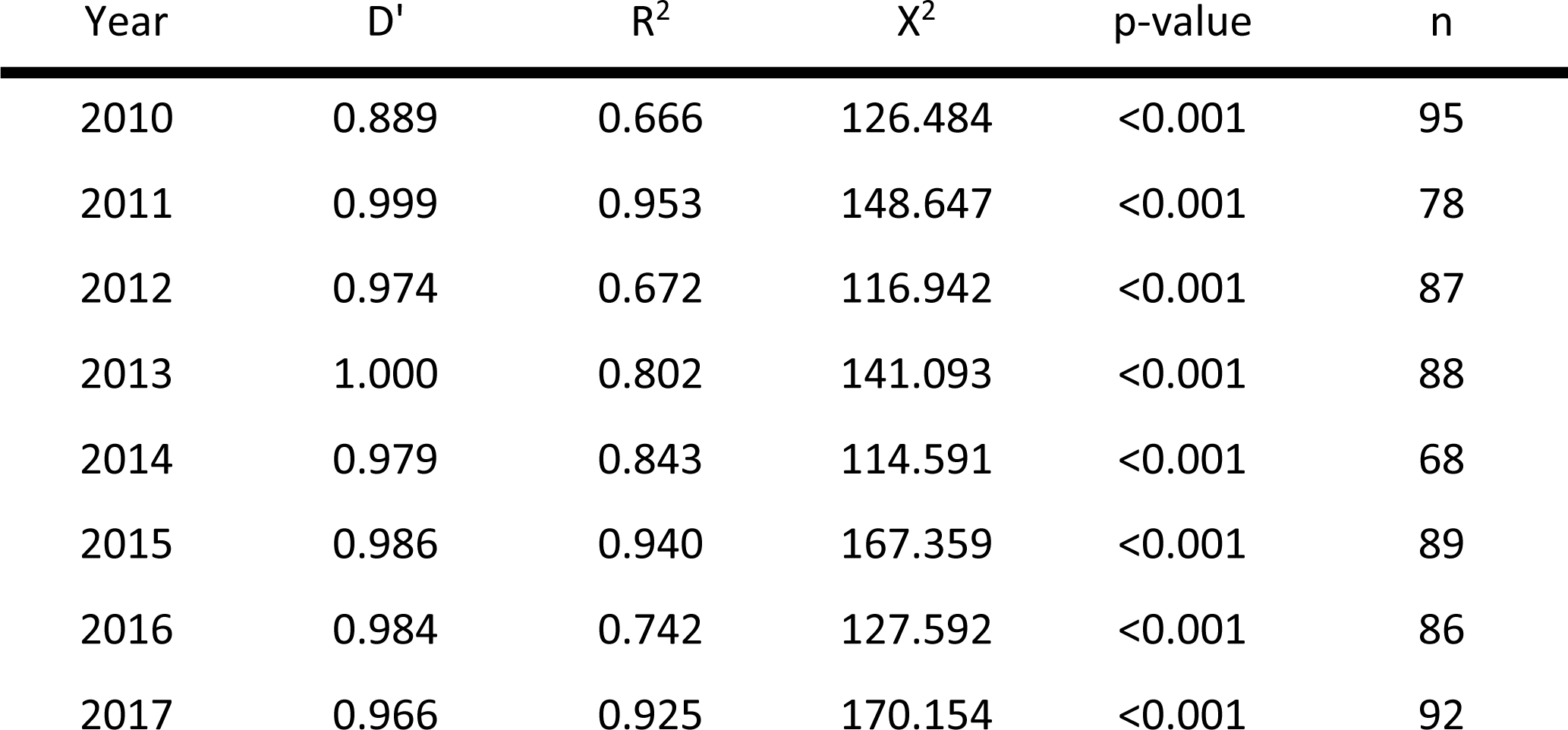
Linkage disequilibrium between V410L and V1016I from 2010 to 2017.

| Year | D' | R<sup>2</sup> | χ<sup>2</sup> | p-value | n |
| --- | --- | --- | --- | --- | --- |
| 2010 | 0.889 | 0.666 | 126.484 | <0.001 | 95 |
| 2011 | 0.999 | 0.953 | 148.647 | <0.001 | 78 |
| 2012 | 0.974 | 0.672 | 116.942 | <0.001 | 87 |
| 2013 | 1.000 | 0.802 | 141.093 | <0.001 | 88 |
| 2014 | 0.979 | 0.843 | 114.591 | <0.001 | 68 |
| 2015 | 0.986 | 0.940 | 167.359 | <0.001 | 89 |
| 2016 | 0.984 | 0.742 | 127.592 | <0.001 | 86 |
| 2017 | 0.966 | 0.925 | 170.154 | <0.001 | 92 |

### Temporal *kdr* Haplotype Frequencies

Prior to the initial use of pyrethroids to control *Ae. aegypti* in Iquitos, most mosquitoes carried the wild type haplotype (Val1016/Phe1534). Immediately after pyrethroid sprays began in 2002 the frequency of the Val1016/Cys1534 haplotype began to rise quickly until it neared fixation in the population in 2006. Notably, the Ile1016/Cys1534 haplotype was not detected until 2010, but then increased in frequency until 2014, when the city stopped spraying pyrethroids and switched to malathion, which has a different mode of action (Fig. 3).

We observed a temporary increase in the wild type haplotype (Val1016/Phe1534) frequency in 2007-2008 (Fig. 3). The 2007-2008 increase was associated with a spatial expansion of the sampling regime that resulted in the southern part of the city being sampled for the first time (Fig. 4c). A large proportion of *Ae. aegypti* in that area of the city carried the wild-type haplotype during this period, but the northern part of the city had a higher proportion of haplotypes containing resistance alleles.

**Figure 4.**
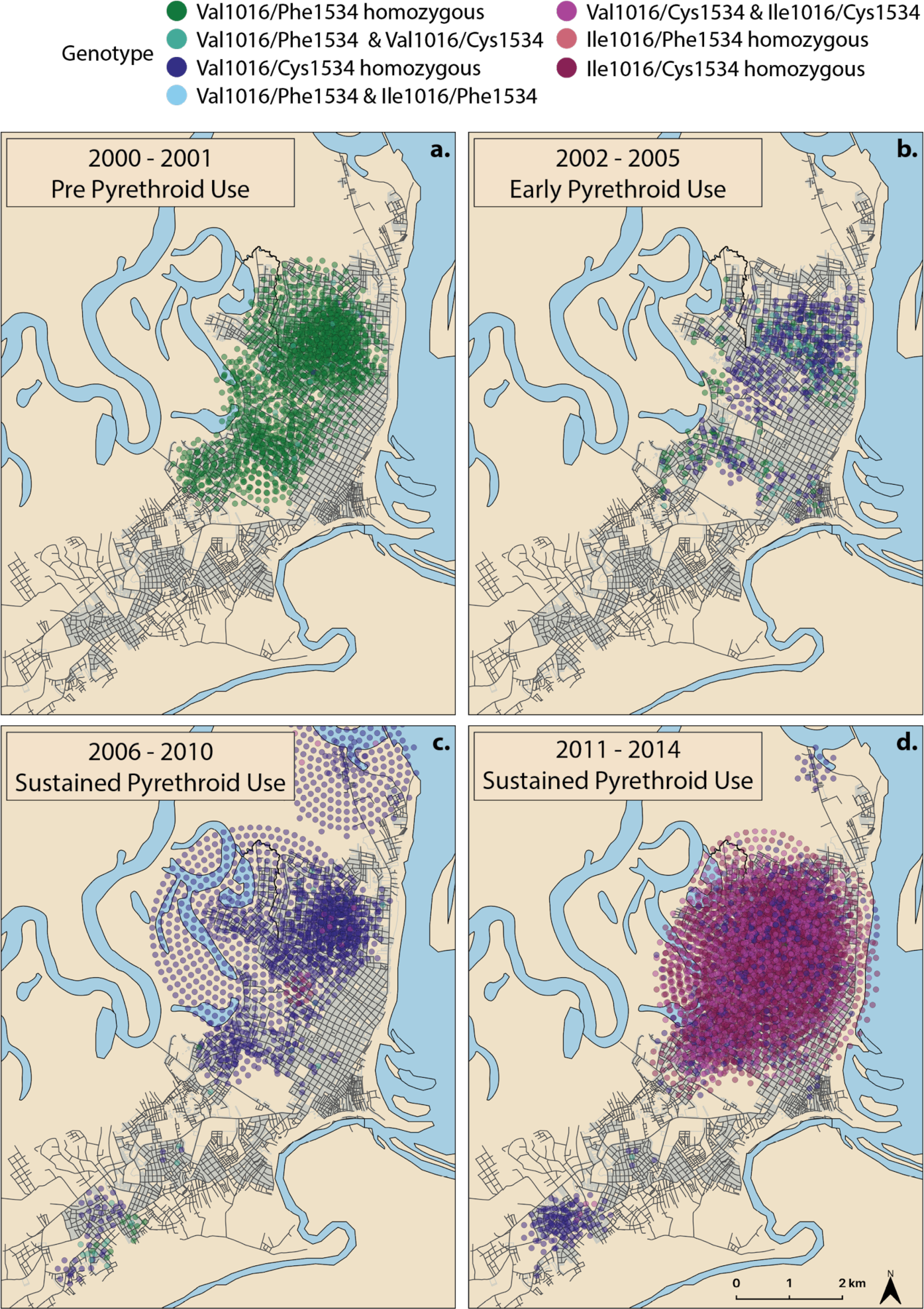
Maps of individual mosquitoes color coded by genotype. Colors similar to haplotypes depicted in Fig. 3. a) 2000 - 2001 pre-pyrethroid use in Iquitos. b) 2002 - 2005 early pyrethroid use. c) 2006 - 2010 sustained pyrethroid use before Ile1016/Cys1534 haplotype emerges. d) 2011 - 2014, sustained pyrethroid use following emergence of Ile1016/Cys1534 haplotype. Some points displaced in concentric circles to show individuals collected from the same location.

During the time period assessed, we observed large and rapid increases in resistance haplotypes even though citywide, low residual, pyrethroid application only occurred on average 3.25 times per year. Considering the inconsistent selection pressure, the mean evolutionary response per mosquito generation (approx. 4-6 weeks) was fairly strong for the resistance alleles at both loci, albeit confidence intervals were wide. Absolute values of the selection coefficients (95% CI) estimated by the WFABC model were 0.313 (0.007, 0.821) and 0.485 (0.145, 0.969) for loci 1016 and 1534, respectively (Fig. 5).

**Figure 5.**
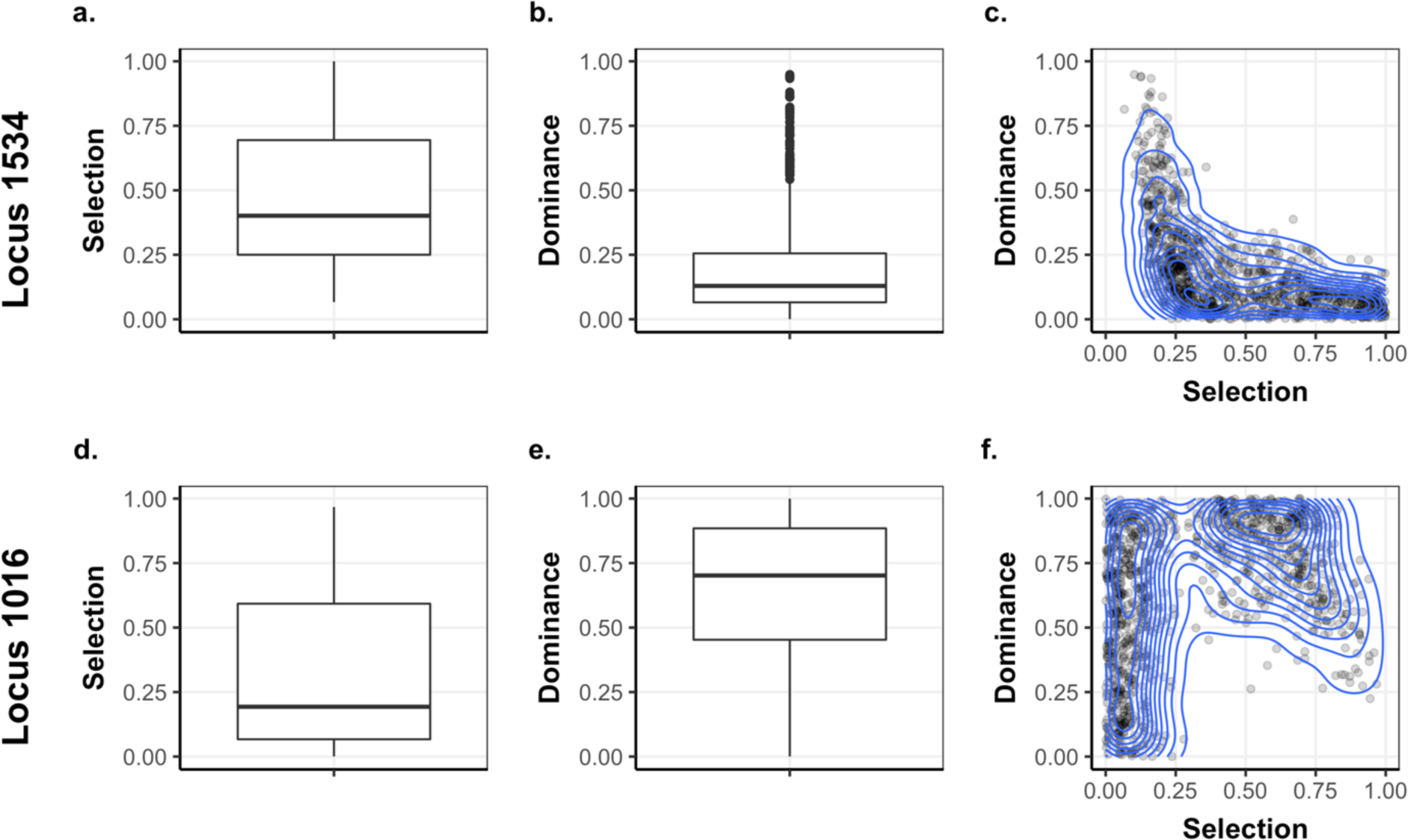
Wright-Fisher Approximate Bayesian Computation (WFABC) results for locus 1534 (top row) and locus 1016 (bottom row). Box plots for the estimated selection coefficients (a, d) and dominance (b, e) from 1000 simulated data sets show the median, 1^st^, and 3^rd^ quartiles, and the whiskers represent the range up to 1.5 times the interquartile range while all outliers are represented by individual points. The points in the 2D plots (c, f) are joint posteriors estimated by the model and the blue curves represent contours of the joint posterior distribution.

Dominance (95% CI) for each resistance allele was estimated by the WFABC model to be 0.635 (0.057, 0.984) and 0.192 (0.007, 0.702), respectively for Ile1016 and Cys1534 (Fig. 5). Recalling that when dominance is zero, the resistance allele is considered dominant, the results indicate that the Ile1016 resistance allele is partially recessive and that the Cys1534 resistance allele is partially dominant to their respective susceptible alleles. Within each locus, the correlation between estimates of the selection coefficient and dominance is illustrated by the two-dimensional joint posterior distribution calculated from the WFABC model (Fig. 5).

Results from our genotype frequency model also indicate that the resistance allele, Cys1534, was partially dominant. Population genetic theory and our deterministic simulations indicate that high values of h cause the resistance allele to take longer to increase from low frequencies, but more quickly approach fixation once at high frequencies, while lower values of h result in a quicker initial increase in frequency, but a longer time to fixation (Fig. 6). The observed dynamics in our system appear to be captured better with smaller values of h in the simulations (Fig. 6 and Suppl Fig. 1). Using pMCMC to estimate parameters of the system, the mean (95% credible interval) for dominance and selection coefficient were *h*=0.184 (0.011,0.571) and *s*=0.188 (0.087,0.284), respectively. The estimated partial dominance of the resistance allele, reflects the behavior of the deterministic simulations, though there is a high degree of uncertainty. Additionally, the estimates for the initial resistance allele frequency and overdispersion parameter were *R_0_* = 0.270 (0.135, 0.450) and *A* = 4.91 (2.97, 7.43). The value for *R_0_* was determined by selecting the resistance allele frequency in October 2002, which is the month that pyrethroid spraying most likely began in earnest in Iquitos. The joint posterior distributions demonstrate correlation between parameter estimates, with a particularly strong correlation between estimates of *h* and *s* and between *R_0_* and *s* (−0.670 and −0.752, respectively), as well as between *h* and *R_0_* (0.536) (Suppl. Fig. 2). For example, without large samples in the first several generations, it may not be possible to distinguish a low initial frequency and large fitness cost from a higher initial frequency but lower cost. We also used the genotype frequency time series samples from pMCMC to construct mean genotype estimates and 95% credible intervals (Suppl. Fig. 3). In later years, these genotype estimates suggest that the susceptible allele is maintained in the population in heterozygotes, which is a result of the low estimates for *h*.

**Figure 6.**
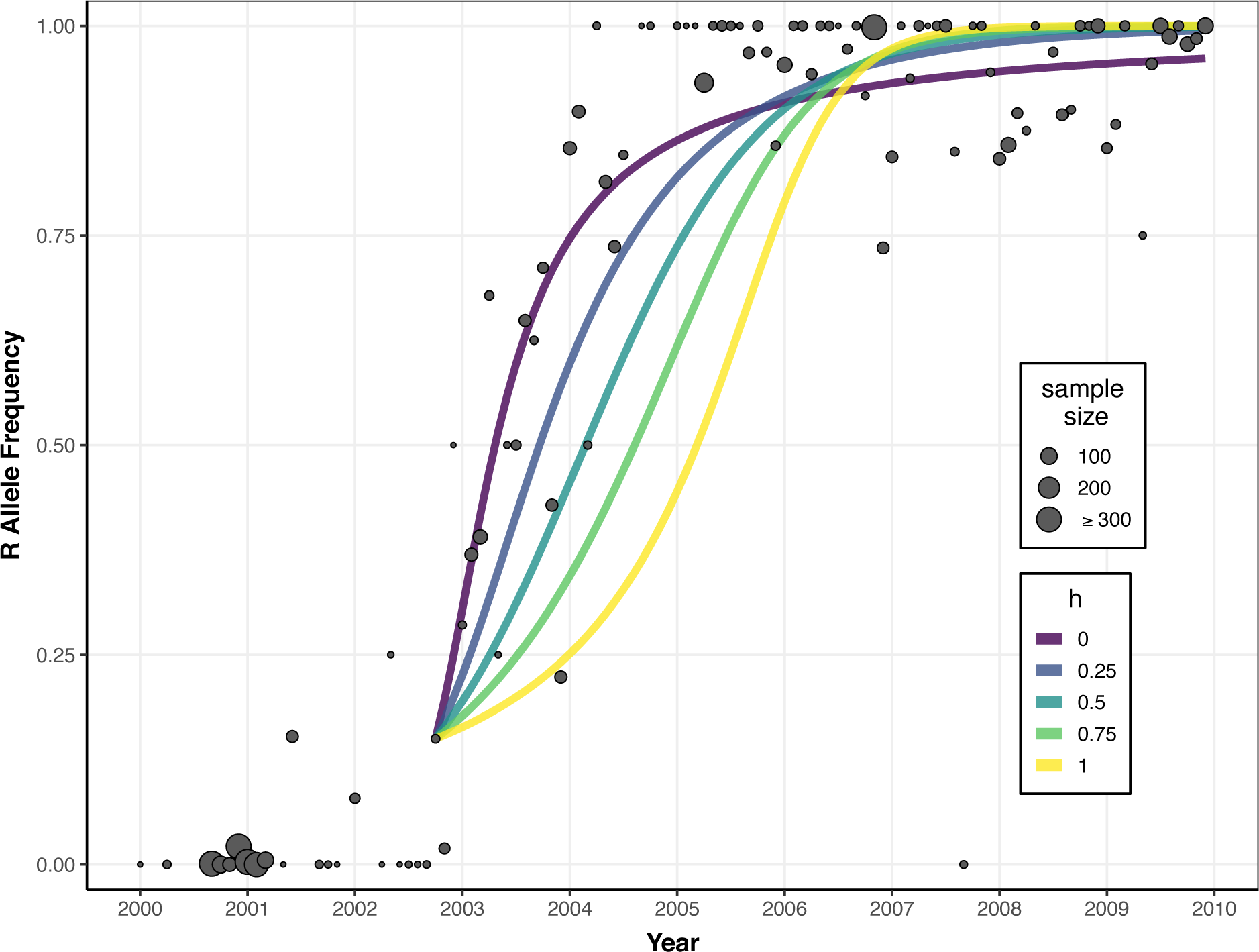
Deterministic model simulations with different dominance parameters. Points indicate empirical 1534 allele (R) frequencies by month, with point size corresponding to the sample size. Model simulations began in October 2002, with initial condition R_0_=0.15 to correspond to the sample of 1 RR, 1 SR, and 8 SS mosquitoes. For each value of *h* (indicated by color), the value of *s* providing the maximum likelihood was used for the simulation.

We noticed that our pMCMC model appeared to predict a greater abundance of heterozygotes than were observed in the empirical data during the period when the Val1016/Cys1534 haplotype was increasing (Suppl. Fig. 3). However, the HWE analysis shows only three months (January 2003, March 2003, December 2003) where our observed genotype frequencies were not in HWE (Suppl. Tbl. 2).

### Spatial *kdr* Haplotype Frequencies

During 2013 and 2014, the haplotype Ile1016/Cys1534 was increasing in frequency in the overall population. We had hypothesized that within each year the frequency of Ile1016/Cys1534 would increase faster in the experimental zone receiving insecticide treatment than in the buffer zone even though there was likely to be movement between the two areas. The sample size from the spray zone for the 2013 suppression experiment was relatively small due to the efficacy of the spraying regime, with 252 mosquitoes analyzed from the buffer zone but only 120 analyzed from the spray zone. After spraying, the resistance allele frequency increased in the spray zone but not the buffer zone (Fig. 7), rising above the buffer zone by 31% in July (p=0.022) and 64% in August (p=0.0068). In 2014, phenotypic resistance was higher than in 2013. A total of 2,178 samples were analyzed for 2014 with a minimum of 52 per group (May, spray zone) and a maximum of 218 per group (October, spray zone) (Fig. 7). Prior to the experimental spray event, the allele frequencies in the spray and buffer zones were not significantly different from each other. During the 6-week intense insecticide selection in the spray zone, the haplotype frequencies of the Ile1016/Cys1534 haplotypes increased in the spray zone relative to the buffer zone by 24% (May, p=0.0011), 19% (July, p=0.0002), and 18% (August, p=0.0091), and finally 11% higher in October (p=0.037) (Fig. 7).

**Figure 7.**
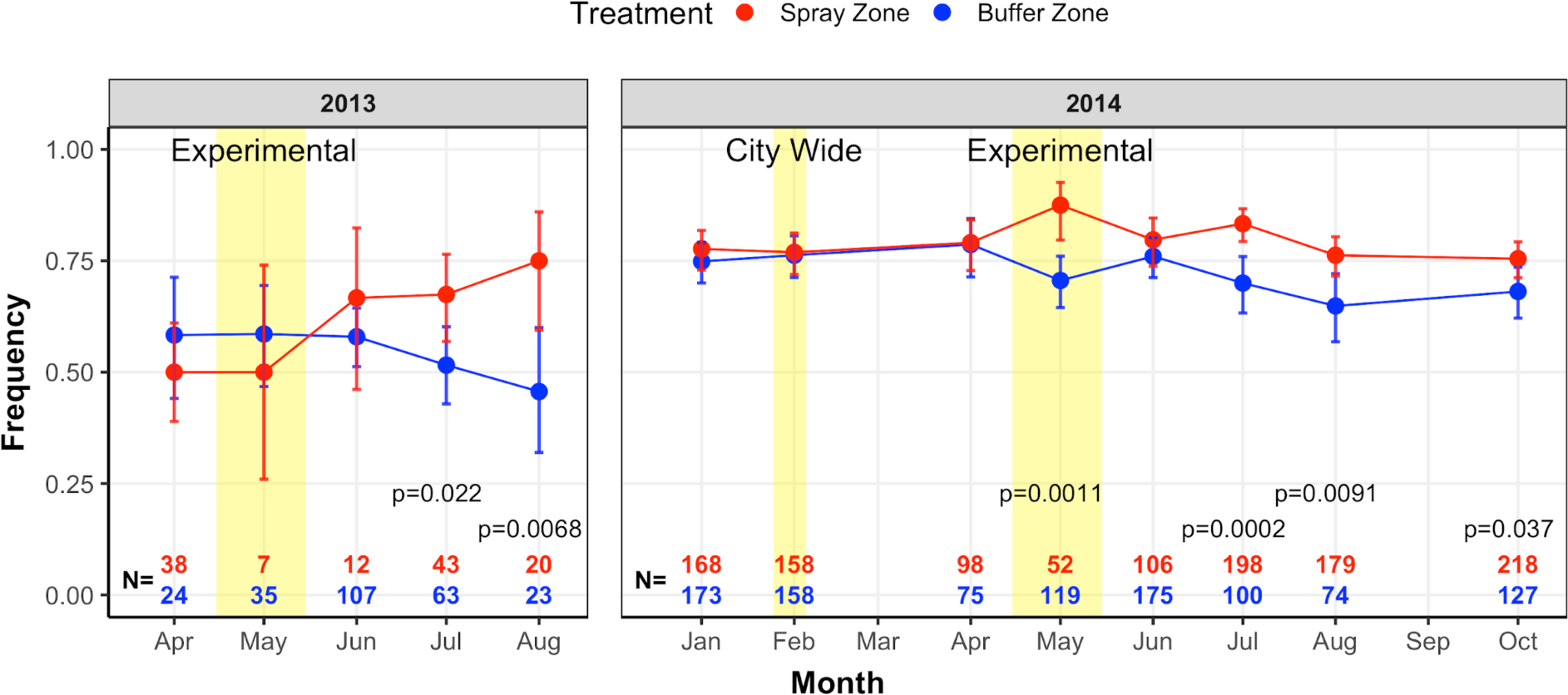
Frequency of Ile1016/Cys1534 haplotype by treatment group across the year 2013 (left) and 2014 (right). A 6-week experimental cypermethrin application, highlighted with yellow blocks, occurred in the spray zone from 29 April - 3 June 2013 and from 28 April - 2 June 2014. A citywide cypermethrin spray in response to a dengue outbreak occurred in February 2014 and is also highlighted with a yellow block. P-values for a test of equality between treatment groups by month were computed from generalized linear models (shown for only for p<0.1).

## Discussion

We have examined the temporal and spatial patterns of *kdr* alleles in *Ae. aegypti* from Iquitos, Peru across an 18-year period, and we identified two episodes of selection for *kdr* haplotypes. We demonstrated that both resistance alleles, Ile1016 and Cys1534, experienced strong selective pressure and that Cys1534 appears to be partially dominant over the wild-type allele. In addition, we detected significant spatial heterogeneity on the scale of a few city blocks, which supports findings reported elsewhere (Deming *et al.,* 2016; Estep *et al.,* 2018; Grossman *et al.,* 2019). Based on the extensive duration and area of sampling, this is the most thorough study of *kdr* population genetics in *Ae. aegypti* to date and it confirms the remarkable capacity for this species to rapidly adapt to insecticidal pressures.

### Haplotype Dynamics

Three *kdr* alleles coding for, Leu410, Ile1016, and Cys1534, that have been shown to contribute to pyrethroid resistance in *Ae. aegypti* in Central and South America were detected in Iquitos. Our data support previous conclusions by Saavedra-Rodriquez *et al*. (2018) who reported evidence for parallel evolution at loci 410 and 1016. The resistance alleles Leu410 and Ile1016 mostly occurred in the same haplotype and rose simultaneously in frequency in the Iquitos population, therefore, only loci 1016 and 1534 were used for haplotype analyses.

In contrast to the haplotype dynamics observed in Vera-Maloof *et al*. (2014) where the allele frequencies of Ile1016 and Cys1534 rose together, in Iquitos, the susceptible wild-type Val1016/Phe1534 haplotype was initially completely replaced by the Val1016/Cys1534 haplotype. Once that haplotype was at or near fixation, the haplotype carrying two resistance alleles, Ile1016/Cys1534 emerged and quickly rose in frequency, replacing the Val1016/Cys1534 haplotype. This indicates that both resistance haplotypes occurred in Iquitos and that there were likely fitness differences between the two haplotypes.

Temporal results indicate that, prior to the municipal citywide spraying of pyrethroids that commenced in 2002, the majority (>99%) of haplotypes in the population were wild-type and <1% of the haplotypes were Val1016/Cys1534 (Fig. 3). Individuals carrying the Val1016/Cys1534 haplotype could either represent standing genetic variation in the population due to mutation/selection balance or to historical selection of the population with DDT, which also interacts with the VGSC protein (Soderlund and Bloomquist 1990; Brengues *et al.,* 2003). However, DDT has not been used in Peru since 1994, and may have last been used in Iquitos in the 1970s (A. Morrison, personal communication), which predates the reintroduction of *Ae. aegypti* to the country. Thus, it is possible that the source population(s) for the reintroduction of the species already contained *kdr* alleles.

### Selection and Dominance

Our data provide field-based evidence that pyrethroid spraying selected for multiple *kdr* haplotypes. This supports previous studies that have demonstrated functional pyrethroid resistance caused by *kdr* mutations in the laboratory (Du *et al.,* 2013, Hirata *et al.,* 2014) and other field-based studies that have reported high frequencies of *kdr* mutations in resistant populations (e.g. Deming *et al.,* 2016; Linss *et al.,* 2014; Li *et al.,* 2015; Brito *et al.,* 2018).

The mosquitoes in Iquitos responded quickly to the new selection pressure from pyrethroids after 2002 when the Val1016/Cys1534 haplotype rapidly rose to high frequencies. The V1016I substitution was not detected in Iquitos until 2010. The later emergence of the resistance allele at the 1016 locus suggests that this genetic mutation was likely not present in the population prior to 2010. In 2010, Iquitos sprayed α-cypermethrin for the first time. The specialized chemistry of this particular formulation may have applied a different selection pressure to the population, resulting in an increase in Ile1016/Cys1534 frequency. Alternatively, the mutation may have arrived in Iquitos via immigration from resistant populations from elsewhere, such as neighboring Brazil where the mutation was first detected in 2006 (Linss *et al.,* 2014). While Iquitos is a fairly isolated city with no roads coming into it from other cities, planes and boats arrive every day and those may facilitate mosquito immigration (Guagliardo *et al*. 2015).

We find evidence of selection pressure in favor of kdr mutations. However, the true selection pressure was likely underestimated because several assumptions of both the WFABC and genotype-frequency models were violated. The selective environment was not constant because the insecticide used, frequency of sprays, and the number of houses sprayed in any given pyrethroid application varied across time and space (Suppl. Tbl. 1). The model also assumed that the loci are independent, but V1016I and F1534C are tightly linked in the genome. Despite the possible underestimation of selection strength, the estimated mean values indicate fairly strong selection pressure for both resistance alleles.

The Cys1534 resistance allele was estimated to be partially dominant to the Phe1534 susceptible allele by both the WFABC model and our genotype-frequency model using pMCMC. While some previous studies have found evidence that *kdr* alleles are recessive, they used laboratory, phenotypic exposure assays to determine inheritance (Saavedra-Rodriguez *et al.,* 2007; Brito *et al.,* 2018). Other investigators have suggested an additive effect with a certain combination of kdr alleles (Ishak *et al.,* 2015; Plernsub *et al.,* 2016). The WFABC method employed in this paper determined inheritance based on the number of generations that an allele existed at low frequencies and did not include phenotypic data (Foll *et al.,* 2014). Phenotypic resistance can be caused by multiple mechanisms within the mosquito, and while the *kdr* mutations can be useful genetic markers for populations under selective pressure, resistance itself is more complex, and involves additional genes and metabolic mechanisms (Smith *et al.,* 2018). In addition, the dominance of an allele can vary depending on environment and genetic background (Bourguet *et al.,* 2000). All of this taken together may explain the estimated differences in dominance between results from this study and previous investigations.

The incomplete dominance estimated here by both the WFABC model and our own population genetics model predicted increases in the rate at which the allele frequencies would rise in the population from small frequencies. This is because most resistance alleles were present as heterozygotes, and higher dominance would give greater selective advantage to the heterozygote compared to the homozygous susceptible under spray conditions. Similarly, when the resistance allele was detected at high frequencies, the susceptible allele was present mostly as heterozygotes and quickly fell out of the population (Conner and Hartl, 2004).

While neither the WFABC model nor our own population genetics model accounted for spatial heterogeneity or migration we know that these phenomena play important roles in *Ae. aegypti* population dynamics. This species has strong spatial structuring, and we know that in the early years of spraying pyrethroids in Iquitos that the populations were significantly reduced. It is possible that these factors are contributing to the rate of resistance evolution rather than the dominance of the Cys1534 resistance allele. Since the dominance estimate for Ile1016 indicated partial recessivity, future investigations may be able to clarify the roles that spatial heterogeneity, migration, and dominance play in the rate of evolution at these two loci. Spatial Heterogeneity

As has been found in previous studies, we were able to detect heterogeneity in *kdr* frequencies at the citywide scale in Iquitos. This is best illustrated by the haplotype frequency data from 2007 and 2008. There appeared to be a decrease in the frequency of resistance alleles in those years (Fig. 3), but this result was likely an artifact of an expanded collection regime and not due to an actual decrease in allele frequency across the city (Fig. 4). Mosquitoes were first sampled in the southern part of the city in 2007. At the time, the northern part of the city had a very high frequency of the Val1016/Cys1534 haplotype, but the south still had a large frequency of the wild type Val1016/Phe1534 haplotype. Phenotypic susceptibility to at least one pyrethroid in this part of the city during this period was confirmed in 2009 by Lenhart *et al*. (2020). By 2011, however, this part of the city also had a high frequency of the Val1016/Cys1534 haplotype, indicating that resistance alleles may have been spreading from the port in the north towards the southern and less populated portion of Iquitos. High frequencies of resistance alleles first occurred in the northern part of the city and spread south over a 10 to 12-year period (approximately, 120 – 144 mosquito generations). It is unknown if these resistance alleles arose independently in Iquitos or were introduced via migration. The Val1016/Cys1534 haplotype already existed within the city prior to widespread pyrethroid selection and it is possible that the Ile1016/Cys1534 haplotype arose via migration but did not reach a detectable frequency until 2010. Future work should explore the demographic history of mosquitoes carrying these haplotypes.

Another indication that spatial heterogeneity exists in Iquitos, can be found in our genotype frequency pMCMC results where we observe fewer heterozygotes than predicted by the model between October 2002 and July 2004 (Suppl. Fig. 3). Although the genotype frequencies are in Hardy-Weinberg equilibrium for 19 out of 22 months (Suppl. Tbl. 2), the deficiency in heterozygotes could be explained by existing genetic structure within the sampled population.

Previous work has indicated that the frequency of resistance alleles within an area can be highly heterogeneous, however, prior studies were unable to determine if the heterogeneity was due to heterogeneity in insecticidal pressures or strong population structure and variation in initial resistance allele frequencies (Linss *et al.,* 2014; Deming *et al.,* 2016). Here, we conducted two suppression experiments over 16 and 44 calendar weeks in years 2013 and 2014, respectively. Significant spatial and temporal heterogeneity in resistance allele frequencies was detected in the 2014 suppression study. While initial resistance allele frequencies were higher that year than the previous year, a large divergence in resistance allele frequency was detected following the experimental 6-week pyrethroid spray treatment, providing further evidence that genetic heterogeneity in *Ae. aegypti* can exist at a very fine spatial scale. In the 2013 experiment, trends indicated that resistance allele frequencies increased in the spray zone compared to the buffer zone. This was likely due to the small sample size available for analysis during that year. Fewer houses were sampled and the spraying regime was much more effective at reducing population size in 2013, likely due to lower levels of resistance compared to the 2014 sampling period.

## Conclusion

This study demonstrated the evolution of two important *kdr* haplotypes in *Ae. aegypti* over the course of 18 years in Iquitos, Peru. We document a strong association between citywide pyrethroid spraying and dramatic increases in the frequency of specific *kdr* haplotypes, including two time periods where dramatic increases in resistance allele frequency were observed in this population. The high estimated selection coefficients for *kdr* haplotypes highlight how quickly the species can adapt to new insecticide pressures even when that pressure is not consistently applied. The rapid speed that resistance allele Cys1534 rose to high frequency may have been aided by the partial dominance this allele exerted over the susceptible allele when under the selection pressure of pyrethroid insecticides. In addition, spatial suppression experiments showed that *kdr* allele frequencies varied not only at the citywide scale, but also on a very fine scale within the city.

Taken together, these results highlight the importance of wide-spread monitoring for insecticide resistance from the beginning of first application. It may be especially important to adjust the spatial scale of monitoring efforts to better capture the potential heterogeneity in allele frequencies that can be found within an area.

## Acknowledgements

We are grateful to the Ministerio de Agricultura y Riego de Peru, Direccion General Forestal y de Fauna Silvestre for permission to conduct these studies under the auspices of Resolución Directoral Nos. 128-2007-Inrena-IFFS-DCB, 415-2009-AG-DGFFS-DGEFFS, 0022-2011-AG-DGFFS-DGEEFFS, 0330-11-AG-DGFFS-DGEFFS, 0306-2013-MINAGRI-DGFFS/DGEFFS. We would like to thank Riddhi Rajyaguru, Laura Welsh, Willy Wei, Shahryar Samir, Destiny Tyson, Lucrecia Vizcaino, Matthew Burrows, Gabriela Vásquez La Torre, and Patricia Barrera for helpful technical and administrative assistance and Brandon Hollingsworth and Sumit Dhole for helpful conversations.

## Supplemental Materials

### Supplemental Methods 1

Detailed description of the genotype-frequency model.

We use a genotype-frequency model to evaluate the spread of the resistance allele at the 1534 locus of the *VGSC*. We assumed the population was well-mixed with random mating and that there were discrete (non-overlapping) generations of mosquitoes, each lasting one month. We also assumed the population was contained, without immigration or emigration of mosquitoes.

We let allele R represent the resistance allele, while allele S represents the wild-type, susceptible allele. The frequencies of each possible genotype were denoted by *X_RR_*, *X_SR_*, and *X_SS_*, with 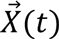 denoting the vector of all three genotype frequencies at generation *t*. For brevity, we omit the vector notation from all vector variables and include a subscript for genotype *i* when referring to a specific element of the vector. The frequency of the R allele in the population is expected to be low until spraying begins because the mutation would likely confer a fitness cost (e.g., proportionately less individuals successfully mate or individuals contribute proportionately fewer offspring) in absence of spraying. For simplicity, we do not consider *de novo* mutations and assume the initial genotype frequencies begin at Hardy-Weinberg equilibrium. Letting *p*(*t*) denote the genotype probabilities of offspring entering generation *t*, we can calculate *p*(*1*) from the starting allele frequency of the resistance allele, *R_0_*:

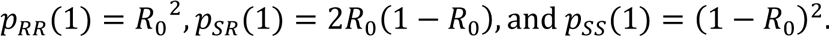

While there is spraying, we assume mosquitoes with an S allele suffer a loss in fitness. Assuming RR is the favored genotype while under selection, SS and SR individuals have a proportionally smaller contribution to the mating pool due to death or fewer offspring. The fitness cost to SS individuals (percent decrease of SS relative to RR) is *s*, and the fitness cost to SR individuals is *h*⋅*s*, where *h* is the dominance of the cost to the S allele.

Using the current frequencies, *X*(*t*), the genotype probabilities of the next generation, *p*(*t*+1), can be calculated based on the probabilities of each of the six possible matings of genotypes. The assumption of a well-mixed and randomly mating population allows the probability of a given mating to be calculated as the product of the pair of genotype frequencies. Each pairing of different genotypes is multiplied by a factor of two to account for the two possible couplings of males and females. The offspring probabilities from each mating are calculated assuming Mendelian inheritance. Fitness costs, as described above, result in a relative reduction in the SR and SS frequencies. The resulting set of difference equations to calculate *p*(*t*+1) from *X*(*t*) (omitting dependence on time for brevity) is:

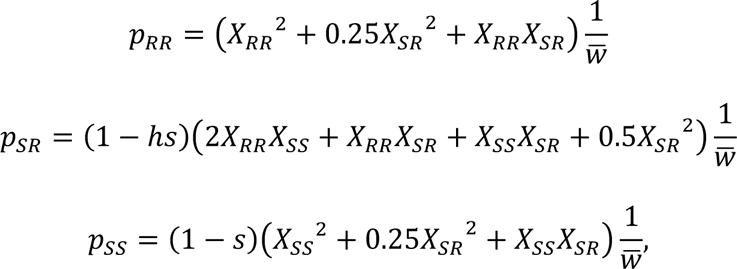

where the mean fitness, 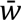, is the sum of the un-normalized frequencies and normalizes so the next generation frequencies sum to 1.

Temporarily ignoring the effects of genetic drift, we can assume the next generation frequencies *X*(*t*+1) are equal to the expected probabilities *p*(*t*+1). By also assuming that the samples are independent and taken at random spatially, maximum likelihood estimation can be used. The data vector from each generation t, *Y*(*t*), consists of counts for each genotype, and we denote the sample size, which varies by generation, by *N_s_*(*t*). By temporarily assuming that samples of individual mosquitoes in a given generation are independent and identically distributed, the sampling distribution of *Y*(*t*) can be approximated by a multinomial distribution with probabilities *X*(*t*) as predicted from simulating the full time series with parameters *θ* = (*s*, *h*, *R*_0_). The likelihood, L, and log-likelihood, *l*, of the data at time t are given by:

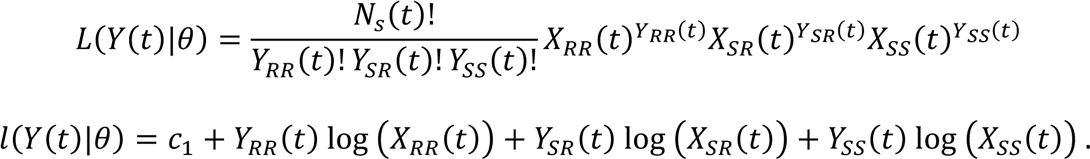

Here, *c_1_* involves terms that do not have a dependence on *X*. Parameter estimates then maximize the log-likelihood of all samples over time for all *T* generations based on the assumption that samples in each generation are independent:

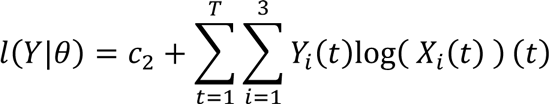

Here, *c_2_* involves terms that do not have a dependence on *X*.

We incorporated stochasticity from genetic drift by assuming a mosquito count equal to the effective population size, *N_e_*, in each generation, assumed to be 500 as in the main text. We let the genotype counts in the next generation, *C*(*t+*1), be distributed multinomially based on the genotype probabilities *p*(*t*). Then we have:

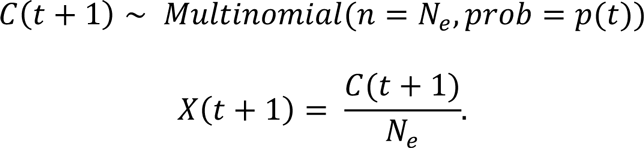

We can additionally loosen our assumption that samples in a given generation are independent and identically distributed. We keep our assumption that the sampling distribution at generation *t* is independent of sampling in previous generations, meaning there is only dependence on the frequencies in that generation, *X*(*t*). We also assume that while sampling removes mosquitoes from the population, the resulting effect on the population genetics is small enough to ignore. To account for the possibility of multiple mosquitoes being sampled from the same house, and heterogeneity in genotype frequencies we employ an overdispersed sampling distribution with added variance compared to a multinomial distribution. Specifically, we employ the Dirichlet-multinomial distribution parameterized by *AX(t)*, where *A* is an overdispersion parameter, which gives the following probability density of *Y* :

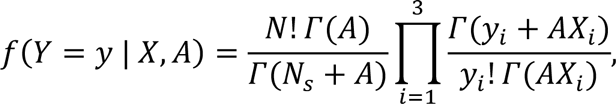

where *Γ*(⋅) is the gamma function. Smaller values of *A* result in higher sample variance, essentially accounting for the possibility of drawing a large fraction of the total samples from few individual houses, which may have mosquito genotype frequencies much different from that of the entire city. To allow for calculation of likelihoods when *X_i_*(*t*) = 0 but *Y_i_*(*t*) > 0, in practice we parameterize the distribution with *AX* + 0.0001. This has a minimal effect on the properties of the distribution but accounts for the possibility of additional sampling error, e.g., if a sample was mislabeled.

The full population genetics model and sampling distribution form a Hidden Markov Model (HMM). We conduct Bayesian inference on the HMM using particle Markov chain Monte Carlo (<u>pMCMC</u>). We implement <u>pMCMC</u> with a multivariate normal proposal distribution and adaptive Metropolis-Hastings acceptance in R (R Core Team 2019) using the package nimble (de Valpine *et al.,* 2017). We used 1000 particles for the particle filter. For parameters in the range [0,1] (*s*, *h*, and *R_0_*), we use uninformative priors of Beta(1,1). For *A*, we use the uninformative prior *A* ∼ *Gamma*(0.01,0.01). To improve the time to convergence, we initialize the parameters using their maximum likelihood estimates. The output of pMCMC is the joint posterior distribution of the parameters along with samples of full time series of genotype frequencies, which we used to construct 95% credible intervals of genotype frequencies.

**Supplemental Table 1.**
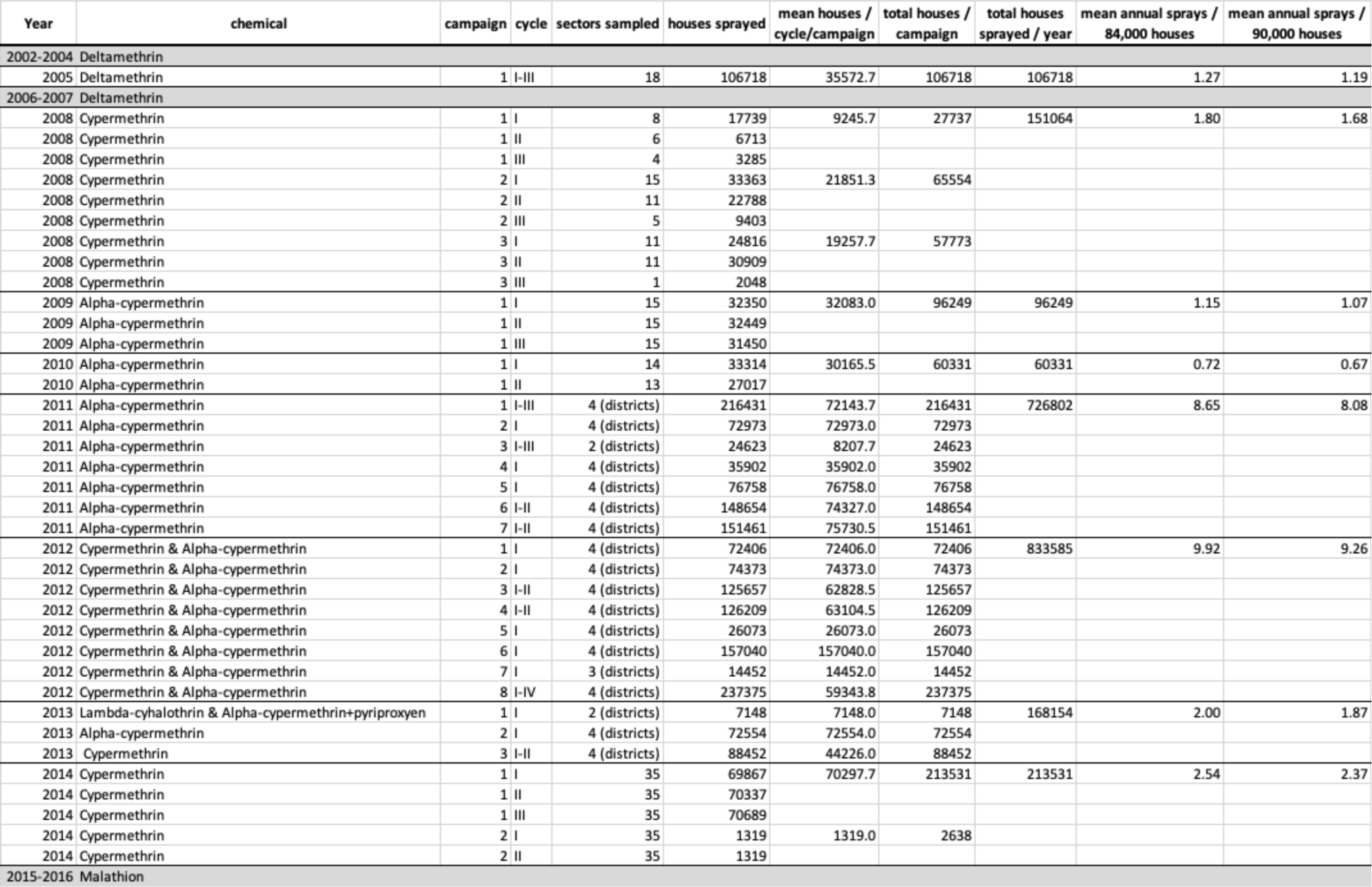
Compiled Ministry of Health insecticide spraying information for Iquitos, Peru from 2000 - 2016.

**Supplemental Table 2.** Hardy-Weinberg equilibrium (HWE) was calculated for select months at locus 1534 using the ‘genetics’ package in R (v. 3.5.0) (Warnes *et al.,* 2019). The Bonferroni significance level is 2.27 x 10^-3^. Months with significant p-values are marked with an asterisk.

| Date | p-value |
| --- | --- |
| 2002-10 | 1.58E-01 |
| 2002-11 | 1.00E+00 |
| 2002-12 | 1.00E+00 |
| 2003-01 | 6.06E-03* |
| 2003-02 | 3.46E-02 |
| 2003-03 | 4.88E-08* |
| 2003-04 | 1.00E+00 |
| 2003-05 | 1.00E+00 |
| 2003-06 | 4.00E-01 |
| 2003-07 | 1.00E+00 |
| 2003-08 | 1.30E-01 |
| 2003-09 | 1.33E-01 |
| 2003-10 | 1.41E-01 |
| 2003-11 | 8.73E-02 |
| 2003-12 | 7.09E-03* |
| 2004-01 | 7.26E-01 |
| 2004-02 | 6.61E-01 |
| 2004-03 | 1.99E-01 |
| 2004-04 | 3.33E-01 |
| 2004-05 | 3.51E-01 |
| 2004-06 | 2.06E-01 |
| 2004-07 | 2.35E-01 |

**Supplemental Figure 1.**
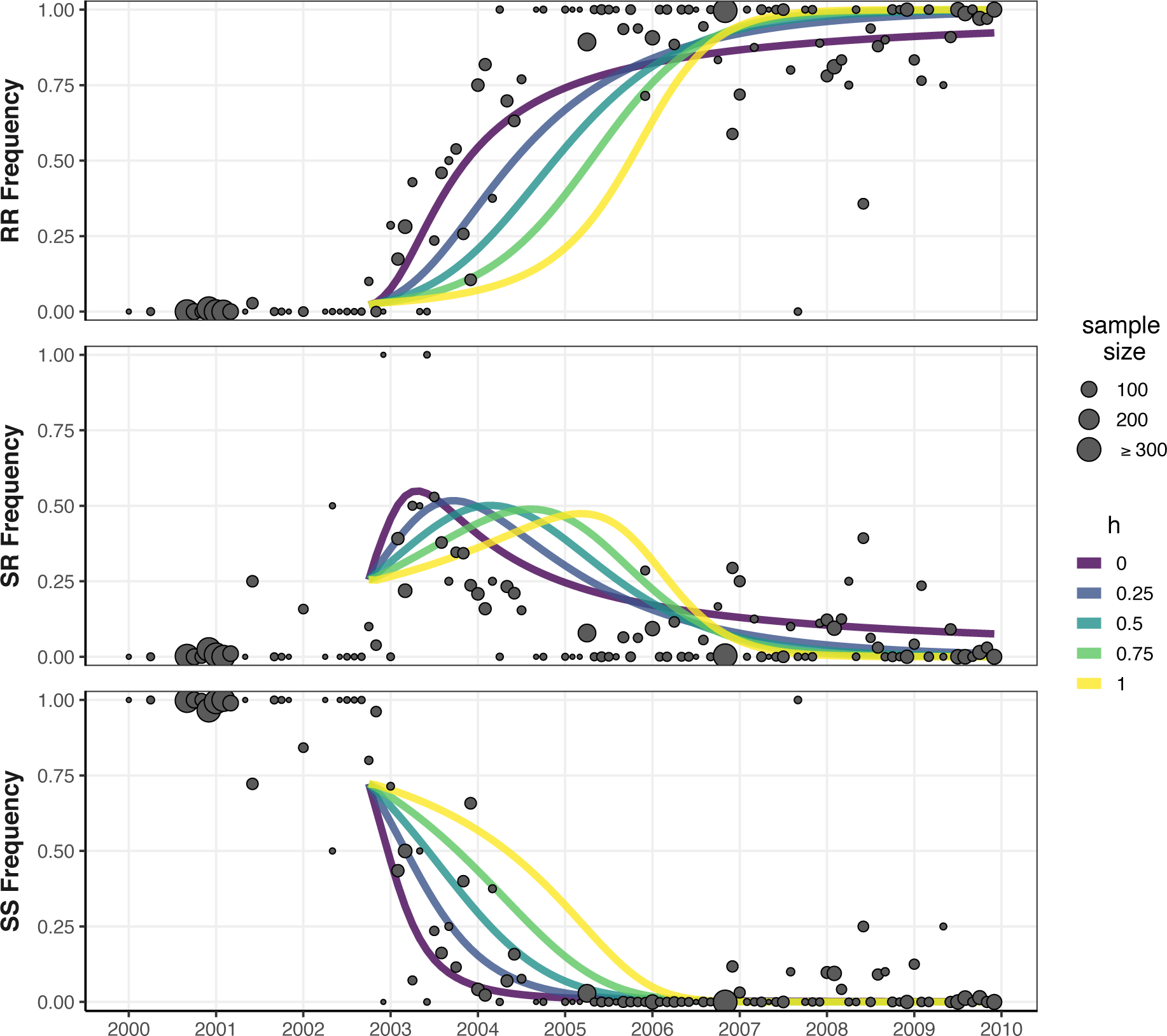
Deterministic model simulations for locus 1534 with different dominance parameters. Points indicate empirical 1534 genotype frequencies, with point size corresponding to the sample size. Model simulations begin in October 2002.

**Supplemental Figure 2.**
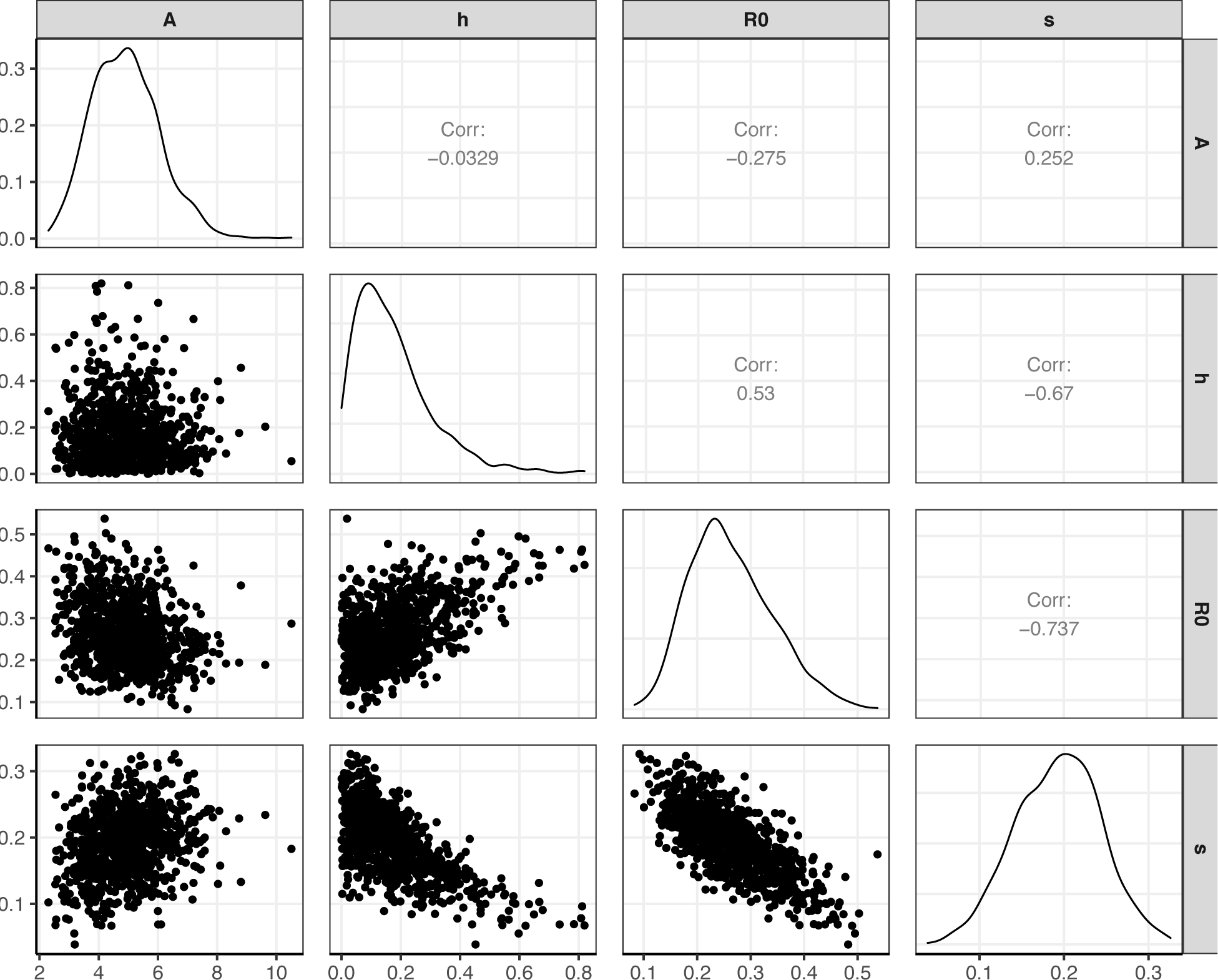
pMCMC pairwise posterior distributions.

**Supplemental Figure 3.**
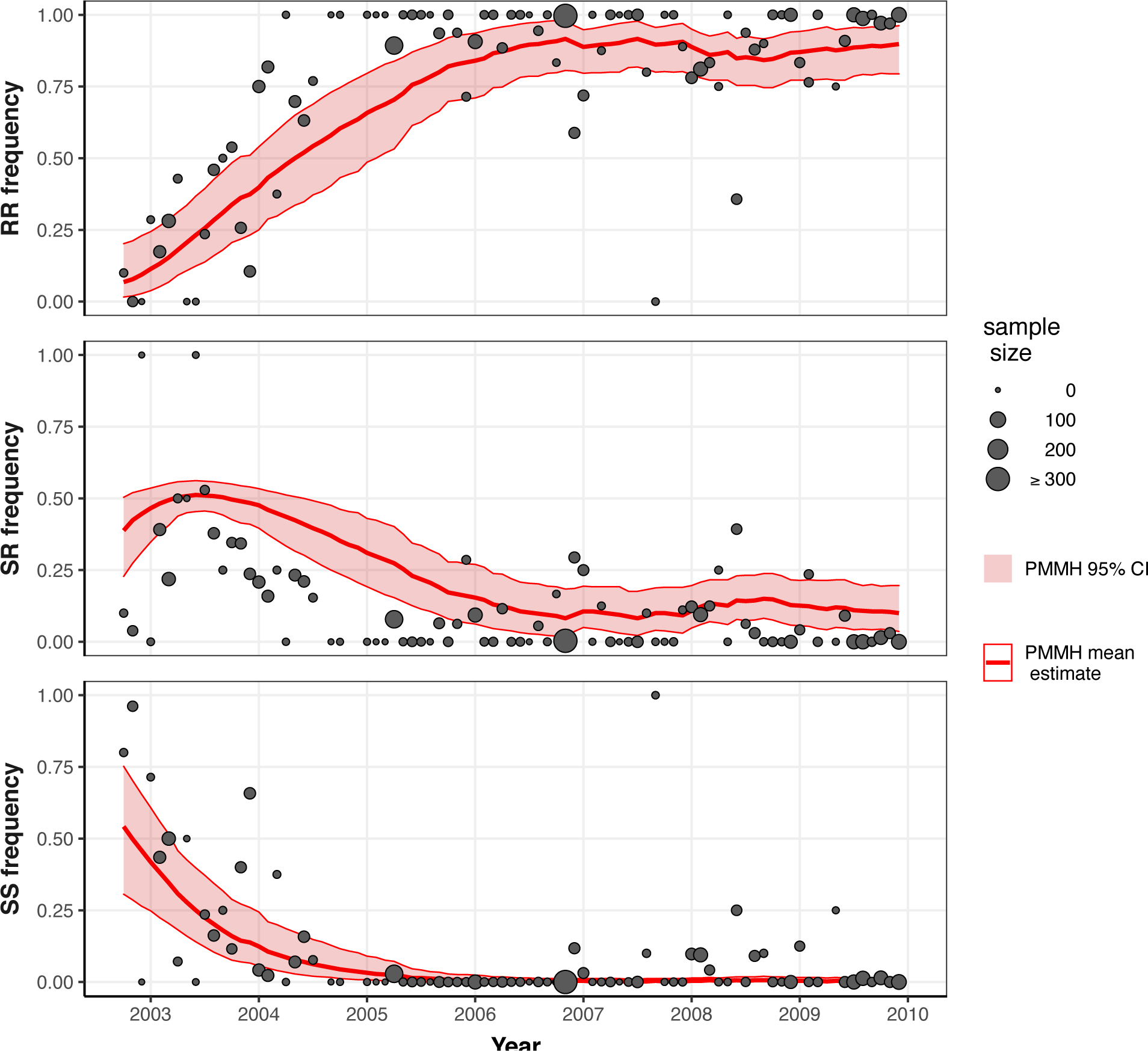
pMCMC genotype frequency estimates and 95% credible intervals. Points indicate empirical 1534 genotype frequencies, with point size corresponding to the sample size. The pMCMC posterior mean genotype frequencies are shown by the red line, and the shaded region shows the 95% credible interval.

## Literature Cited

[dataset]Authors; Year; Dataset title; Data repository or archive; Version (if any); Persistent identifier (e.g. DOI). *To be completed upon acceptance*.

Bharati, M., Saha D. (2018). Multiple insecticide resistance mechanisms in primary dengue vector, *Aedes aegypti* (Linn.) from dengue endemic districts of sub-Himalayan West Bengal, India. PLoS ONE, 13(9), e0203207.

Bhatt, S., Gething, P.W., Brady, O.J., Messina, J.P., Farlow, A.W., Moyes, C.L., Drake, J.M., Brownstein, J.S., Hoen, A.G., Sankoh, O., et al. (2013). The global distribution and burden of dengue. Nature, 496, 504–507.

Brengues, C., Hawkes, N.J., Chandre, F., McCarroll, L., Duchon, S., Guillet, P., Manguin, S., Morgan, J.C., Hemingway, J. (2003). Pyrethroid and DDT cross-resistance in *Aedes aegypti* is correlated with novel mutations in the voltage-gated sodium channel gene. Med Vet Entomol, 17(1), 87–94.

Brito, L.P., Carrara, L., de Freitas, R.M., Lima, J.B.P., Martins, A.J. (2018). Levels of resistance to pyrethroid among distinct *kdr* alleles in *Aedes aegypti* laboratory lines and frequency of *kdr* alleles in 27 natural populations from Rio de Janeiro, Brazil. BioMed Res Int, 2018, 1–10.

Canadian Center for DNA Barcoding. (2020, July 1). CCDB – DNA Extraction. http://ccdb.ca/site/wp-content/uploads/2016/09/CCDB_DNA_Extraction.pdf

Carvalho, D.O., McKemey, A.R., Garziera, L., Lacroix, R., Donnelly, C.A., Alphey, L., Malavasi, A., Capurro, M.L. (2015). Suppression of a field population of *Aedes aegypti* in Brazil by sustained release of transgenic male mosquitoes. PLoS Negl Trop Dis, 9(7), e0003864.

Cavany, S. M., España, G., Lloyd, A. L., Waller, L. A., Kitron, U., Astete, H., Elson, W. H., Vazquez-Prokopec, G. M., Scott, T. W., Morrison, A. C., Reiner, R. C., Jr, & Perkins, T. A. (2020). Optimizing the deployment of ultra-low volume and targeted indoor residual spraying for dengue outbreak response. PLoS Computational Biology, 16(4), e1007743.

Clark, G.G., Seda, H., Gubler, D.J. (1994). Use of the “CDC backpack aspirator” for surveillance of *Aedes aegypti* in San Juan, Puerto Rico. J Am Mosq Control Assoc, 10(1), 119–24.

Conner, J.K., D. L. Hartl. (2004). A primer of ecological genetics. Sinauer Associates Incorporated.

Cosme, L.V. et al. (2020). Evolution of Kdr Haplotypes in Worldwide Populations of Aedes Aegypti: Independent Origins of the F1534C Kdr Mutation. PLoS Neglected Tropical Diseases, 14(4), e0008219–18.

Cromwell, E.A., Stoddard, S.T., Barker, C.M., Van Rie, A., Messer, W.B., Meshnick, S.R., Morrison, A.C., Scott, T.W. (2017). The relationship between entomological indicators of Aedes aegypti abundance and dengue virus infection. PLoS Neglected Tropical Diseases, 11(3), e0005429.

de Valpine, P. et al. (2017). Programming with Models: Writing Statistical Algorithms for General Model Structures with NIMBLE. Journal of Computational and Graphical Statistics, 26(2), 403–413.

Deming, R., Manrique-Saide, P., Barreiro, A.M., Cardena, E.U.K., Che-Mendoza, A., Jones, B., Liebman, K., Vizcaino, L., Vazquez-Prokopec, G., Lenhart, A. (2016). Spatial variation of insecticide resistance in the dengue vector *Aedes aegypti* presents unique vector control challenges. Parasite Vector, 9(67), 1–10.

Dong, K., Du, Y., Rinkevich, F., Nomura, Y., Xu, P., Wang, L., Silver, K., Zhorov, B.S. (2014). Molecular biology of insect sodium channels and pyrethroid resistance. Insect Biochem Molec, 50, 1–17.

Du, Y., Nomura, Y., Zhorov, B., Dong, K. (2016). Sodium channel mutations and pyrethroid resistance in *Aedes aegypti*. Insects, 7(4), 60.

Estep, A.S., Sanscrainte, N.D., Waits, C.M., Bernard, S.J., Lloyd, A.M., Lucas, K.J., et al. (2018). Quantification of permethrin resistance and *kdr* alleles in Florida strains of *Aedes aegypti* (L.) and *Aedes albopictus* (Skuse). PLoS Negl Trop Dis, 12(10), e0006544.

Fan, Y., et al. (2020). Evidence for Both Sequential Mutations and Recombination in the Evolution of Kdr Alleles in Aedes Aegypti. PLoS Neglected Tropical Diseases, 14(4), e0008154–22.

Flores-Suarez, A.E., Ponce-Garcia, G., Lopez-Monroy, B., Villanueva-Segura, O.K., Rodriguez-Sanchez, I.P., Arredondo-Jimenez, J.I., Manrique-Saide, P. (2016). Current Status of the Insecticide Resistance in Aedes aegypti (Diptera: Culicidae) from Mexico. Insecticides Resistance doi:10.5772/61526.

Foll, M., Shim, H., Jensen, J.D. (2014). WFABC: a Wright-Fisher ABC-based approach for inferring effective population sizes and selection coefficients from time-sampled data. Mol Ecol Resour, 15(1), 87–98.

GBD 2017 Causes of Death Collaborators. Global, regional, and national age-sex-specific mortality for 282 causes of death in 195 countries and territories, 1980-2017: a systematic analysis for the Global Burden of Disease Study 2017. Lancet. 2018 Nov 10;392(10159):1736–1788. doi: 10.1016/S0140-6736(18)32203-7. Epub 2018 Nov 8. Erratum in: Lancet. 2019 Jun 22;393(10190):e44. Erratum in: *Lancet*. 2018 Nov 17;392(10160):2170. PMID: 30496103; PMCID: PMC6227606.

Getis, A., Morrison, A.C., Gray, K., Scott, T.W. (2003). Characteristics of the spatial pattern of the dengue vector, *Aedes aegypti*, in Iquitos, Peru. Am. J. Trop. Med. Hyg, 69(5), 494–505.

Grossman, M.K., Rodriguez, J., Barreiro, A.M., Lenhart, A., Manrique-Saide, P., Vazquez-Prokopec, G.M. (2019). Fine-scale spatial and temporal dynamics of *kdr* haplotypes in Aedes aegypti from Mexico. Parasite Vector, 12(20), 1–12.

Guagliardo, S.A., Morrison, A.C., Barboza, J.L., Requena, E., Astete, H., Vazquez-Prokopec, G., Kitron, U. (2015). River boats contribute to the regional spread of the Dengue vector *Aedes aegypti* in the Peruvian Amazon. PLoS Negl Trop Dis, 9(4), e0003648.

Gunning, C.E., Okamoto, K.W., Astete, H., Vasquez, G.M., Erhardt, E., Del Aguila, C., Pinedo, R., Cardenas, R., Pacheco, C., Chalco, E., et al. (2018). Efficacy of *Aedes aegypti* control by indoor Ultra Low Volume (ULV) insecticide spraying in Iquitos, Peru. PLoS Negl Trop Dis, 12(4), e0006378.

Haddi, K., Tomé, H.V.V., Du, Y., Valbon, W.R., Nomura, Y., Martins, G.F., Dong, K., Oliveira, E.U. (2017). Detection of a new pyrethroid resistance mutation (V410L) in the sodium channel of Aedes aegypti: a potential challenge for mosquito control. Sci Rep, 7, 46549.

Harris, A.F., McKemey, A.R., Nimmo, D., Curtis, Z., Black, I., Morgan, S.A., Oviedo, M.N., Lacroix, R., Naish, N., Morrison, N.I., et al. (2012). Successful suppression of a field mosquito population by sustained release of engineered male mosquitoes. Nature Biotechnol, 30(9), 828–830.

Hartl, DL. (2000). A primer of population genetics. Edition 3. Massachusetts (MA): Sinauer Associates, Inc.

Hoffmann, A.A., Montgomery, B.L., Popovici, J., Iturbe-Ormaetxe, I., Johnson, P.H., Muzzi, F., Greenfield, M., Durkan, M., Leong, Y.S., Dong, Y., et al. (2011). Successful establishment of Wolbachia in Aedes populations to suppress dengue transmission. Nature, 476(7361), 454–457.

Instituto Nacional de Estadistica e Informatica. (2012). Perú: Estimaciones y Proyecciones de Población Total por Sexo de las Principales Ciudades, 2000--2015. Boletín Especial N° 23.

Ishak, I.H., Jaal, Z., Ranson, H., Wondji, C.S. (2015). Contrasting patterns of insecticide resistance and knockdown resistance (*kdr*) in the dengue vectors *Aedes aegypti* and *Aedes albopictus* from Malaysia. Parasite Vector, 8(181), 1–13.

Kamgang, B., Yougang, A.P., Tchoupo, M., Riveron, J.M., Wondji, C. (2017). Temporal distribution and insecticide resistance profile of two major arbovirus vectors *Aedes aegypti* and *Aedes albopictus* in Yaoundé, the capital city of Cameroon. Parasite Vector, 10(469), 1–9.

LaCon, G., Morrison, A.C., Astete, H., Stoddard, S.T., Paz-Soldan, V.A., Elder, J.P., Halsey, E.S., Scott, T.W., Kitron, U., Vazquez-Prokopec, G.M. (2014). Shifting patterns of *Aedes aegypti* fine scale spatial clustering in Iquitos, Peru. PLoS Negl Trop Dis, 8(8), e3038.

Lacroix, R., McKemey, A.R., Raduan, N., Wee, L.K., Ming, W.H., Ney, T.G., Rahidah, A.A.S., Salman, S., Subramaniam, S., Nordin, O., et al. (2012). Open field release of genetically engineered sterile male *Aedes aegypti* in Malaysia. PLoS One, 7(8), e42771.

Lenhart, A., Morrison, A. C., Paz-Soldan, V. A., Forshey, B. M., Cordova-Lopez, J. J., Astete, H., Elder, J. P., Sihuincha, M., Gotlieb, E. E., Halsey, E. S., Kochel, T. J., Scott, T. W., Alexander, N., & McCall, P. J. (2020). The impact of insecticide treated curtains on dengue virus transmission: A cluster randomized trial in Iquitos, Peru. PLoS Neglected Tropical Diseases, 14(4), e0008097.

Lenth, R. (2020). emmeans: Estimated Marginal Means, aka Least-Squares Means. R package version 1.5.2-1. https://CRAN.R-project.org/package=emmeans

Li, C., Kaufman, P.E., Xue, R., Zhao, M., Wang, G., Yan, T., Guo, X., Zhang, Y., Dong, Y., Xing, D., et al. (2015). Relationship between insecticide resistance and *kdr* mutations in the dengue vector *Aedes aegypti* in Southern China. Parasite Vector, 8(325), 1–9.

Linss, J.G.B., Brito, L.P., Garcia, G.A., Araki, A.S., Bruno, R.V., Lima, J.B.P., Valle, D., Martins, A.J. (2014). Distribution and dissemination of the Val1016Ile and Phe1534Cys *Kdr* mutations in *Aedes aegypti* Brazilian natural populations. Parasite Vector, 7(25), 1–11.

Magori, K., Legros, M., Puente, M.E., Focks, D.A., Scott, T.W., Lloyd, A.L., Gould, F. (2009). Skeeter Buster: A stochastic, spatially explicit modeling tool for studying *Aedes aegypti* population replacement and population suppression strategies. PLoS Negl Trop Dis, 3(9), e508.

Morrison, A.C., Gray, K., Getis, A., Astete, H., Sihuincha, M., Focks, D., Watts, D., Stancil, J.D., Olson, J.G., Blair, P., Scott, T.W. (2004). Temporal and geographic patterns of *Aedes aegypti* (Diptera: Culicidae) production in Iquitos, Peru. J. Med. Entomol, 41(6), 1123–1142.

Morrison, A.C., Astete, H., Chapilliquen, F., Ramirez-Prada, C., Diaz, G., Getis, A., Gray, K., Scott, T.W. (2004). Evaluation of a sampling methodology for rapid assessment of Aedes aegypti infestation levels in Iquitos, Peru. Journal of medical entomology, 41, 502–510.

Morrison, A.C., Sihuincha, M., Stancil, J.D., Zamora, E., Astete, H., Olson, J.G., Vidal-Ore, C., Scott, T.W. (2006). Aedes aegypti (Diptera: Culicidae) production from non-residential sites in the Amazonian city of Iquitos, Peru. Annals of tropical medicine and parasitology, 100 Suppl 1:S73–S86.

Morrison, A.C., Forshey, B.M., Notyce, D., Astete, H., Lopez, V., Rocha, C., Carrion, R., Carey, C., Eza, D., Montgomery, J.M., Kochel, T.J. (2008). Venezuelan equine encephalitis virus in Iquitos, Peru: urban transmission of a sylvatic strain. PLoS neglected tropical diseases, 2, e349.

Murcia, O., Henriquez, B., Castro, A., Koo, S., Young, J., Márquez, R., Pérez, D., Cáceres, L., Valderrama, A. (2019). Presence of the point mutations Val1016Gly in the voltage-gated sodium channel detected in a single mosquito from Panama. Parasite Vector, 12(62), 1–7.

Nguyen, T.H., Nguyen, H.L., Nguyen, T.Y., Vu, S.N., Tran, N.D., Le, T.N., Vien, Q.M., Buh, T.C., Le, H.T., Kutcher, S., et al. (2015). Field evaluation of the establishment potential of wMelPop Wolbachia in Australia and Vietnam for dengue control. Parasite Vector, 8(563), 1–14.

Nguyen, T.K.L., Nguyen, T.H.B., Nguyen, T.H.N., Nguyen, T.H., Nguyen, H.H. (2018). Two novel mutations in the voltage-gated sodium channel associated with knockdown resistance (*kdr*) in the dengue vector *Aedes aegypti* Vietnam. J Vector Ecol, 43(1), 184–189.

Plernsub, S., Saingamsook, J., Yanola, J., Lumjuan, N., Tippawangkosol, P., Sukontason, K., Walton, C., Somboon, P. (2016). Additive effect of knockdown resistance mutations, S989P, V1016G and F1534C, in a heterozygous genotype conferring pyrethroid resistance in *Aedes aegypti* in Thailand. Parasite Vector, 9(417), 1–7.

Phillips, I., Need, J., Escamilla, J., Colán, E., Sánchez, S., Rodríguez, M., Vásquez, L., Seminario, J., Betz, T., da Rosa, A.T. (1992). First Documented Outbreak of Dengue in the Peruvian Amazon Region. Bulletin of Pan American Health Organization, 26(3), 201–207.

R Project for Statistical Computing. © The R Foundation. (2020). R Foundation for Statistical Computing, Vienna, Austria. Available at: https://www.R-project.org/.

Reiner, R.C., Jr, Stoddard, S.T., Vazquez-Prokopec, G.M., Astete, H., Perkins, T. A., Sihuincha, M., Stancil, J.D., Smith, D.L., Kochel, T. J., Halsey, E.S., Kitron, U., Morrison, A.C., and Sott, T.W. (2019). Estimating the impact of city-wide Aedes aegypti population control: An observational study in Iquitos, Peru. PLoS Negl. Trop. Dis, 13, e0007255.

Saarman, N.P., Gloria-Soria, A., Anderson, E.C., Evans, B.R., Pless, E., Cosme, L.V., Gonzalez-Acosta, C., Kamgang, B., Wesson, D.M., Powell, J.R. (2017). Effective population sizes of a major vector of human diseases, *Aedes aegypti*. Evol Appl, 10(10), 1031–1039.

Saavedra-Rodriguez, K., Maloof, F.V., Campbell, C.L., Garcia-Rejon, J., Lenhart, A., Penilla, P., Rodriguez, A., Sandoval, A.A., Flores, A.E., Ponce, G., et al. (2018). Parallel evolution of *vgsc* mutations at domains IS6, IIS6 and IIIS6 in pyrethroid resistant *Aedes aegypti* from Mexico. Sci Rep, 8(6747), 1–9.

Saavedra-Rodriguez, K., Urdaneta-Marquez, L., Rajatileka, S., Moulton, M., Flores, A.E., Fernandez-Salas, I., Bisset, J., Rodriguez, M., Mccall, P.J., Donnelly, M.J., et al. (2007). A mutation in the voltage-gated sodium channel gene associated with pyrethroid resistance in Latin American *Aedes aegypti*. Insect Molecular Biology, 16(6), 785–798.

Schneider, J.R., Morrison, A.C., Astete, H., Scott, T.W., Wilson, M.L. (2004). Adult size and distribution of Aedes aegypti (Diptera: Culicidae) associated with larval habitats in Iquitos, Peru. Journal of medical entomology, 41, 634–642.

Smith, L.B., Kasai, S., Scott, J.G. (2016). Pyrethroid resistance in *Aedes aegypti* and *Aedes albopictus*: Important mosquito vectors of human diseases. Pestic Biochem Phys, 133, 1–12.

Smith, L.B., Sears, C., Sun, H., Mertz, R.W., Kasai, S., Scott, J.G. (2019). CYP-mediated resistance and cross-resistance to pyrethroids and organophosphates in *Aedes aegypti* in the presence and absence of *kdr*. Pestic Biochem Phys, 160, 119–126.

Soderlund, D.M., Bloomquist, J.R. (1990). Molecular Mechanisms of Insecticide Resistance. Pesticide Resistance in Arthropods. Springer. p. 21–56.

Thomas, S.J., Yoon, I.K. (2019). A review of Dengvaxia®: development to deployment. Hum Vacc Immunother, 15(10), 2295–2314.

Tun-Lin, W., Lenhart, A., Nam, V.S., Rebollar-Tellez, E., Morrison, A.C., Barbazan, P., Cote, M., Midega, J., Sanchez, F., Manrique-Saide, P., Kroeger, A., Nathan, M.B., Meheus, F., Petzold, M. (2009). Reducing costs and operational constraints of dengue vector control by targeting productive breeding places: a multi-country non-inferiority cluster randomized trial. Tropical medicine & international health, 14, 1143–1153.

Vera-Maloof, F.Z., Saavedra-Rodriguez, K., Elizondo-Quiroga, A.E., Lozano-Fuentes, S., Black, W.C., IV. (2015). Coevolution of the Ile1,016 and Cys1,534 mutations in the voltage gated sodium channel gene of *Aedes aegypti* in Mexico. PLoS Negl Trop Dis, 9(12), e0004263.

Warnes, G., with contributions from Gorjanc G, Leisch F, Man M. (2019). genetics: Population Genetics. R package version 1.3.8.1.2. Available at: https://CRAN.R-project.org/package=genetics.

Watts, D.M., Porter, K.R., Putvatana, P., Vasquez, B., Calampa, C., Hayes, C.G., Halstead, S.B. (1999). Failure of secondary infection with American genotype dengue 2 to cause dengue hemorrhagic fever. The Lancet, 354(9188), 1431–1434.

World Health Organization. (2020, July 1). Vector-borne Diseases. https://www.who.int/news-room/fact-sheets/detail/vector-borne-diseases.

WHO Expert Committee on Vector Biology and Control. (1992). Vector resistance to pesticides: fifteenth report of the WHO Expert Committee on Vector Biology and Control. World Health Organization, 1992.

Yanola, J., Somboon, P., Walton, C., Nachaiwieng, W., Somwang, P., Prapanthadara, L. (2011). High-throughput assays for detection of the F1534C mutation in the voltage-gated sodium channel gene in permethrin-resistant *Aedes aegypti* and the distribution of this mutation throughout Thailand. Trop Med Int Health, 16(4), 501–509.

